# Neural Mechanisms of Feedback Processing and Regulation Recalibration during Neurofeedback Training

**DOI:** 10.1101/2024.08.19.608543

**Authors:** Gustavo S. P. Pamplona, Jana Zweerings, Cindy S. Lor, Lindsay deErney, Erik Roecher, Arezoo Taebi, Lydia Hellrung, Kaoru Amano, Dustin Scheinost, Florian Krause, Monica D. Rosenberg, Silvio Ionta, Silvia Brem, Erno J. Hermans, Klaus Mathiak, Frank Scharnowski

## Abstract

The acquisition of new skills is facilitated by providing individuals with feedback that reflects their performance. This process creates a closed loop that involves feedback processing and regulation recalibration to promote effective training. Functional magnetic resonance imaging (fMRI)-based neurofeedback is unique in applying this principle by delivering direct feedback on the self-regulation of brain activity. Understanding how feedback-driven learning occurs requires examining how feedback is evaluated and how regulation adjusts in response to feedback signals. In this pre-registered mega-analysis, we re-analyzed data from eight intermittent fMRI neurofeedback studies (N = 153 individuals) to investigate brain regions where activity and connectivity are linked to feedback processing and regulation recalibration (i.e., regulation after feedback) during training. We harmonized feedback scores presented during training in these studies and computed their linear associations with brain activity and connectivity using parametric general linear model analyses. We observed that, during feedback processing, feedback scores were positively associated with (1) activity in the reward system, dorsal attention network, default mode network, and cerebellum; and with (2) reward system-related connectivity within the salience network. During regulation recalibration, no significant associations were observed between feedback scores and either activity or associative learning-related connectivity. Our results suggest that neurofeedback is processed in the reward system, supporting the theory that reinforcement learning shapes this form of brain training. In addition, the involvement of large-scale networks in feedback processing, continuously transitioning between evaluating external feedback and internally assessing the adopted cognitive state, suggests that higher-level processing is integral to this type of learning. Our findings highlight the pivotal role of performance-related feedback as a driving force in learning, potentially extending beyond neurofeedback training to other feedback-based processes.

**Key Points:** We conducted a pre-registered mega-analysis integrating data from eight fMRI neurofeedback studies to examine feedback processing and regulation recalibration during neurofeedback training.

During feedback processing, feedback was associated with activity in the reward system, dorsal attention network, default mode network, and cerebellum; as well as with reward system-related connectivity within the salience network.

We found no positive results during regulation blocks; however, additional analyses suggest that recalibration may have already occurred during feedback presentation.

## 1. Introduction

Feedback can facilitate and aid the acquisition of new skills by providing individuals with information about their performance. The information has to be evaluated and compared to the previous action in order to improve task performance. This process creates a closed-loop system, where feedback is constantly updated and individuals adjust mental strategies accordingly. In order to maximize positive and minimize negative feedback, individuals learn by adapting their mental strategies, which involves processing feedback information (requiring emotional valuation and memory), and the motivation for and planning of new mental strategy. This learning process is known as reinforcement learning.

Reinforcement learning is usually associated with the modulation of dopaminergic activity in the reward system, a brain network comprising limbic regions (midbrain, ventral and dorsal striatum, orbitofrontal cortex, ventromedial prefrontal cortex, amygdala, thalamus), as well as the anterior and posterior cingulate, insular cortex, and inferior frontal gyrus (O’Doherty, 2004; Schultz et al., 1997; Tricomi and DePasque, 2016; Tsukamoto et al., 2006). Several functional imaging studies have shown greater activity in several of these regions in response to positive feedback (Elliott et al., 2003; Marco-Pallarés et al., 2007; Nieuwenhuis et al., 2005b, 2005a). The limbic system is thought to have an important role in feedback processing due to its involvement in motivation and assigning value to information (Haber, 2011). Notably, the nucleus accumbens (NAcc) is a key integrative region for feedback processing, motivation, and learning, and is thought to modulate behavior according to a goal (Goto and Grace, 2005), being connected to the amygdala, the hippocampus (Goto and Grace, 2008), the ventral tegmental area (Camara et al., 2009; Knutson and Gibbs, 2007), as well as other limbic regions and cortical regions. After receiving feedback on task performance, the individual can reframe cognitive strategies to improve performance. The dlPFC plays a critical role in associative learning due to its engagement in working memory, attentional switching, and response selection (Niendam et al., 2012). Top-down attentional control is primarily associated with the posterior parietal cortex (and intraparietal sulcus) (Corbetta and Shulman, 2002; Green and McDonald, 2008). Alongside parietal regions, the anterior cingulate cortex and NAcc have been suggested to be involved in behavior adaptation (Holroyd and Coles, 2002).

Feedback is essential for learning, but when it comes to training the brain directly, we do not have conscious access to our own brain activity. Neurofeedback overcomes this shortcoming by converting brain activity into sensory feedback (e.g., a visual thermometer). Using neurofeedback, participants can learn to modulate their own brain activity voluntarily, which can lead to behavioral changes (Sitaram et al., 2017; Weiskopf et al., 2004). Neurofeedback experiments entail a distinctive form of learning, wherein the objective is to improve the self-regulation of a particular neural signal. Neurofeedback learning is thought to be driven by reinforcement learning, whereby neural states become more probable when they are associated with performance-related rewards (Lubianiker et al., 2022; Shibata et al., 2019; Sitaram et al., 2017). In this context, effective feedback processing represents a crucial factor in reinforcement learning. Several other factors make neurofeedback a worthwhile paradigm to investigate the neural mechanisms of feedback processing and regulation recalibration. Often, neurofeedback experiments provide graded feedback rather than binary feedback, which can be studied using parametric general linear models (GLM) (Radua et al., 2018). Neurofeedback experiments also often employ the acquisition with whole-brain coverage. Finally, neurofeedback studies using intermittent feedback, as opposed to continuous feedback, separate feedback presentation and periods of self-regulation into temporally distinct blocks, allowing feedback evaluation to be examined without interference from self-regulation (Johnson et al., 2012; Lubianiker et al., 2022). Although several cognitive theories have been proposed to elucidate the mechanisms underlying learning in neurofeedback experiments (Lubianiker et al., 2022; Sitaram et al., 2017), and the focus of some studies on brain responses due to feedback during neurofeedback training (Dewiputri et al., 2021; Emmert et al., 2016; Hinterberger et al., 2005; Mathiak et al., 2015; Radua et al., 2018; Shibata et al., 2019), the relationship between feedback scores and brain changes during feedback and regulation blocks in the context of neurofeedback training remains poorly understood.

Here, our goal was to capitalize on existing neurofeedback studies to investigate the brain correlates of feedback processing and regulation recalibration, i.e., adapting or reinforcing a regulation strategy depending on the feedback received. More specifically, we reanalyzed a dataset of 153 participants from eight fMRI-based intermittent neurofeedback studies using parametric GLMs. Pooling data from several neurofeedback studies allows for examining generalizable neural mechanisms while minimizing study-specific effects (e.g., feedback representation, directionality of regulation, and target brain regions). We only included studies using intermittent feedback to make sure that the feedback processing and self-regulation are temporally not overlapping.

We hypothesized that, in the context of neurofeedback training, (1) during feedback presentation blocks, performance-related feedback scores are positively associated with activity in reward-related brain regions (ventral tegmental area, NAcc, ventral striatum, anterior cingulate cortex, anterior insular cortex), and (2) with connectivity between those regions. We also hypothesized that (3) during regulation blocks, performance-related feedback scores on the previous trial are negatively associated with activity in brain regions related to neurofeedback control (dorsolateral prefrontal cortex, posterior parietal cortex, lateral occipital cortex, thalamus) and (4) with connectivity between those regions. The negative association posited in hypotheses 3 and 4 would indicate that a reduction in the level of performance-related feedback received would necessitate greater effort for the recalibration of the regulatory strategy, resulting in an increase in brain activity/connectivity.

Testing these hypotheses will contribute to our understanding of how the brain processes feedback and how regulation is recalibrated after feedback presentation in neurofeedback training. The insights we gain from this study might also generalize to other feedback-based learning contexts that depend on reward processing and regulation recalibration. Thus, our findings may inform neuroscientific theories of learning and future brain-based interventions to optimize learning.

## 2. Materials and Methods

The hypotheses and the analyses were preregistered prior to conducting the study: osf.io/bzweg/

### 2.1. Data collection procedure

The present study is a mega-analysis (Costafreda, 2009), which involved systematically searching the literature for relevant studies, requesting data sharing from the authors, and analyzing the raw data from neurofeedback training using a different approach than that employed in the offline analyses of the original studies.

First, we systematically searched for articles whose data could potentially be included in our mega-analysis study using the Scopus systematic search (www.scopus.com). The full procedure for the systematic search is described in the first section of the Supp. Material. Inclusion criteria were (1) original research on neurofeedback, (2) studies using fMRI as the acquisition technique, (3) paradigms using intermittent (rather than continuous) neurofeedback, and (4) published as peer-reviewed scientific articles. The search identified seven studies that could be included in our mega-analysis. Two additional studies were included that were not identified in the search but met the inclusion criteria for this analysis (Amano et al., 2016; Zweerings et al., 2019). We contacted all authors via e-mail and inquired about data sharing of raw anatomical and functional imaging data, performance-related feedback scores for each feedback block, and basic information about the experimental design (e.g., block onsets and durations). We also requested minimal descriptive demographic information on gender and age. Of the nine selected studies, we obtained consent and collected data from eight fMRI-neurofeedback studies (Amano et al., 2016; Hellrung et al., 2018; Keller et al., 2021; Krause et al., 2021; Pamplona et al., 2020; Scheinost et al., 2020; Zweerings et al., 2020, 2019). This study protocol was approved by the ethics committee of the University of Vienna (EK 00621).

### 2.2. Participants

Relevant demographic information about each study is described in Table 1. Irrespective of the original design, only data from healthy participants who received intermittent and veridical feedback were considered. Participants who received sham feedback (Scheinost et al., 2020) were not included in this study. Due to technical issues in the data sharing process, data from some participants could not be recovered (four from Zweerings et al., 2019, and one from Hellrung et al., 2018) resulting in a data set of 153 participants.

**Table 1.** Demographic details of each included study (studies ordered by sample size after study-specific exclusion criteria).

| Study | Handedness | Number of groups in the original study | Head motion as an exclusion criterion | Number of participants* | Mean age | Number of females* | Sham feedback group |
| --- | --- | --- | --- | --- | --- | --- | --- |
| Keller et al., 2021 | Right | 2 (patients and HC) | Yes. Two HC participants removed from the analysis | 37 | 32.3 ± 2.1 | 15 | No |
| Zweerings et al., 2019 | Right | 2 (schizophrenia patients and HC) | No | 35 | 34.4 ± 12.1 | 16 | No |
| Zweerings et al., 2020 | Right | 3 (2 PTSD in a cross-over design, 1 HC) | No | 21 | 44.1 ± 10.9 | 9 | No |
| Hellrung et al., 2018 | Right | 3 (1 continuous, 1 intermittent feedback, 1 no feedback) | Yes. Scan-to-scan movements of more than 3 mm | 18 | 27.8 ± 3.8 | 0 | No |
| Pamplona et al., 2020 | Right | 2 (1 NF, 1 behavioral test-retest only) | No | 15 | 26.9 ± 3.8 | 10 | No |
| Amano et al., 2016 (some additional details of this study can be found in Cortese et al., 2021) | Not collected | 2 (1 NF, 1 no feedback) | No | 12 | 21.2 ± 1.8 | 1 | No |
| Scheinost et al., 2020 | Right | 2 (NF and sham feedback) | Yes | 10 | 21.5 ± 2.6 | 10 | Yes, yoked feedback |
| Krause et al., 2021 | Right:8, Left:1, Ambidexterity:1 | 1 | No | 10 | 25.8 ± 6.2 | 5 | No |

*Note.* \* After exclusion, considering only the relevant group for the present study. HC = healthy controls; PTSD = post-traumatic stress disorder; NF = neurofeedback.

### 2.3. fMRI acquisition parameters and experimental design of the studies

A summary of the fMRI acquisition parameters is shown in the supplementary material (Table S1). For all studies, data acquisition was performed using 3T MRI scanners, echo-planar sequences and axial slice orientation. A summary of the experimental design for each study can also be found in the supplementary material (Table S2).

### 2.4. Data Analysis

#### 2.4.1. Harmonization of feedback values

We investigated linear associations between performance-related feedback during neurofeedback training and estimates of brain activity and connectivity in parametric analyses (i.e., over a range of feedback scores), rather than in a categorical analysis (i.e., positive versus negative feedback). Feedback scores from each dataset were harmonized across studies before computing these associations.

In the first step, we converted feedback scores into numerical values. Three studies presented a two-digit representation proportional to the previous block of self-regulation performance (Keller et al., 2021; Zweerings et al., 2020, 2019). Accordingly, no numerical transformation was conducted for these studies. For the other five studies, feedback was presented graphically in the form of thermometers (Hellrung et al., 2018; Pamplona et al., 2020), concentric discs (Amano et al., 2016; Krause et al., 2021), or a speedometer (Scheinost et al., 2020). We converted the feedback scores of all datasets to decimal numbers in the range 0-1, proportional to the level of steps compared to the maximum and minimum possible representations provided after the regulation block. Datasets from three studies (Hellrung et al., 2018; Krause et al., 2021; Zweerings et al., 2019) combined up- and down-regulation trials during neurofeedback training. To reflect feedback representation of success and failure independent of directionality, we inverted the scale for feedback presentation blocks following down-regulation (i.e., 0 and 1 represented the upper and lower feedback limits, respectively). For one study, the requested direction of feedback representation was rightward (Scheinost et al., 2020), so 0 and 1 represented the leftmost and rightmost feedback limits, respectively.

For some datasets, the converted feedback values showed a highly non-uniform distribution that could affect the linear regression of the parametric analysis. In extreme cases, a high number of feedback scores near the upper or lower limits would lead, in practice, to a categorical analysis (most commonly, near the upper and lower limits). Therefore, we applied the following iterative procedure to remove highly frequent range-specific values. First, for each individual, we calculated the histogram of feedback scores with *h* bins, in which *h* is the number of possible subject-specific graphical feedback steps between the minimum and maximum feedback scores received by the subject during training (Hellrung et al., 2018; Keller et al., 2021; Pamplona et al., 2020; Zweerings et al., 2020, 2019). In some studies, continuous graphical feedback steps were presented, and *h* was determined by subtracting for each subject the minimum feedback score from the maximum and dividing the result by 100 (Amano et al., 2016; Krause et al., 2021; Scheinost et al., 2020). We then calculated the mean and standard deviation (SD) of the counts across the bins. If the bin with the highest count of feedback scores was above the threshold of the mean value plus three SD, one score from a randomly selected feedback presentation block belonging to that bin was marked as censored. Then, the mean and SD of the counts across bins were recalculated, and the procedure was repeated until the bin with the highest count was not higher than the threshold. Finally, scores from feedback presentation blocks marked as censored were removed from the analyses. The final mean percentage (± SD) of censored feedback scores across individuals in each dataset was 30.0 ± 15.1% for Keller et al., 2021; 40.3 ± 14.7% for Zweerings et al., 2019; 7.1 ± 10.6% for Zweerings et al., 2020; 1.2 ± 2.5% for Hellrung et al., 2018; zero for Pamplona et al., 2020; 81.5 ± 6.5% for Amano et al., 2016; and 4.2 ± 4.4% for Scheinost et al., 2020; 0.1 ± 0.2 for Krause et al., 2021. The final mean number (± SD) of maintained feedback scores across individuals in each dataset was 50.4 ± 10.9 for Keller et al., 2021; 41.6 ± 13.0 for Zweerings et al., 2019; 16.7 ± 2.0 for Zweerings et al., 2020; 23.7 ± 0.6 for Hellrung et al., 2018, 40 for Pamplona et al., 2020; 98.6 ± 34.9 for Amano et al., 2016; 11.5 ± 0.5 for Scheinost et al., 2020; 375.5 ± 8.4 for Krause et al., 2021.

#### 2.4.2. fMRI preprocessing

We applied the same preprocessing pipeline to the requested raw data to minimize variance across studies that could arise from disparities in MRI acquisition parameters and preprocessing. All fMRI data were preprocessed prior to statistical mapping using MATLAB (version R2022b, www.mathworks.com) and the Statistical Parametric Mapping toolbox (SPM12, Wellcome Department of Imaging Neuroscience, University College London, UK). We first slice-time corrected functional MRI data (according to the slice acquisition order of the dataset) using the middle slice as the reference. From the resulting images, we estimated the translation and rotation parameters of head motion, and resliced the images to a created mean image using a fourth-degree B-spline interpolation. We then coregistered the anatomical image to this mean image using a rigid-body model. The coregistered anatomical image was used to generate a deformation field to normalize the resulting anatomical and functional images of all subjects according to the standard Montreal Neurological Institute (MNI) stereotactic space with a resolution of 2 x 2 x 2 mm³. Finally, we performed spatial smoothing using a Gaussian kernel of 8 mm³ at full-width half maximum (FWHM).

#### 2.4.3. First-level analysis of the association between performance-related feedback and estimates of activity/connectivity

We performed voxel-wise mass-univariate general linear model (GLM) analyses using SPM12 to determine regions whose activity/connectivity are parametrically associated with feedback values during feedback presentation blocks as well as during the subsequent regulation blocks. One GLM was specified separately for each model to test for distinct hypotheses.

Prior to GLM analyses, we concatenated the runs of each individual to center feedback scores subject-wise rather than run-wise, to increase the number of blocks for parametric analysis and to extract single time series for connectivity analyses. This step was done using the SPM ‘spm_fmri_concatenate’ function (github.com/spm/spm12/blob/master/spm_fmri_concatenate.m), which includes run-specific regressors to remove run effects (henceforth, run-effect regressors) and corrects the high-pass filter and non-sphericity calculations.

#### Model 1

Model 1 investigated the association between performance-related feedback and activity during feedback blocks (Hypothesis 1). As indicated in the preregistration, for some datasets, the last feedback block of each run was not included in the regressor because this block was at the end of the training run and its delayed hemodynamic response could not be determined. For each individual, we constructed two regressors, one for feedback presentation blocks and one for regulation blocks, as boxcar functions using study-specific onsets and durations. The (unmodulated) regressor representing feedback presentation was orthogonalized with a parametrically modulated regressor representing the performance-dependent feedback score presented in each block.

#### Model 2

Model 2 was designed to investigate brain regions whose connectivity with a reward-related region, the NAcc, is associated with the level of performance-related feedback during feedback blocks (Hypothesis 2). The estimated connectivity during the feedback presentation was computed using the psycho-physiological interaction (PPI) analysis (O’Reilly et al., 2012). Similar to Model 1, the last feedback block of each run was not considered in some datasets. We used the IBASPM 71 atlas (Alemán-Gómez et al., 2006) within the WFU PickAtlas toolbox (Maldjian et al., 2003, www.nitrc.org/projects/wfu_pickatlas/) to create a binary mask of the bilateral NAcc. We then extracted the averaged time course within the NAcc across concatenated runs and computed the element-by-element product of this time course and the “task” regressor. This task regressor was an HRF-convolved boxcar function constructed from the modulated feedback presentation blocks. Variance explained by the six head motion parameters, run-effect regressors, and regressors of no interest was removed. Finally, we estimated the first-level betas voxel-wise for each GLM based on the PPI regressors.

#### Model 3

Model 3 investigated the association between performance-related feedback and activity during regulation blocks (Hypothesis 3). The first regulation block of each run was not included in the model, because there was no feedback presentation block prior to the first regulation block. As announced in the preregistration, we constructed four regressors representing feedback presentation blocks, baseline blocks, the first four seconds of the regulation blocks (referred to here as “onset of regulation”), and the remaining regulation block. We included only the beginning of the regulation blocks in the model, as we assumed that modulation would be stronger during the initial phase of a new strategy, compared to its maintenance over the remainder of the block. Although regulation recalibration driven by feedback may occur later in the block, we focus here exclusively on the initial phase due to our assumption that feedback-driven recalibration happens primarily at the beginning of the regulation block followed by a lower degree of feedback-driven maintenance of activity. Methodologically, adopting a short and consistent duration for modeling the initial recalibration phase helps control for the wide variation in regulation block durations across studies, which range from 6 seconds (e.g., Amano et al., 2016) to 180 seconds (e.g., Scheinost et al., 2020; see Table S2). Additionally, the (unmodulated) regressor representing the onset of regulation was orthogonalized with a parametrically modulated regressor corresponding to performance-related feedback score presented in each previous feedback block.

#### Model 4

Model 4 was designed to investigate brain regions whose connectivity with the dorsolateral prefrontal cortex (dlPFC) is associated with performance-related feedback during the onset of regulation blocks (Hypothesis 4). The estimated connectivity during the onset these blocks was computed using PPI analysis. Similar to Model 3, the first regulation block in each run was not considered. The left dlPFC was selected as the ROI due to its established link to neurofeedback control (Emmert et al., 2016; Sitaram et al., 2017). To define this ROI, we first downloaded a meta-analytic map from Neurosynth (https://neurosynth.org/) using the term “dlpfc”. Then, using the MarsBaR toolbox (<u>marsbar.sourceforge.net</u>), we created a 6-mm-radius spherical ROI centered on the MNI coordinate peak (-46, 38, 30) from the meta-analytic map to represent the dlPFC. We then extracted the averaged time course within the dlPFC across concatenated runs and computed the element-by-element product of this time course and the “task” regressor. This task regressor was an HRF-convolved boxcar function based on the modulated onset of regulation blocks. Variance explained by the six head motion parameters, run-effect regressors, and regressors of no interest was removed. Finally, we estimated the first-level betas voxel-wise for each GLM based on the PPI regressors.

All parametric modulators were specified as first order (linear association). In addition to unmodulated, modulated, and run-effect regressors, the six head motion parameters estimated in the preprocessing step were added to the first-level design matrices. For six datasets (Hellrung et al., 2018; Keller et al., 2021; Krause et al., 2021; Pamplona et al., 2020; Zweerings et al., 2020, 2019), we applied a high-pass filter with a 128-s cut-off to remove the low-frequency signal. Due to the long periods between feedback presentation blocks, a high-pass filter with a cut-off of half the run duration (Nurmi et al., 2018) was applied to the remaining two datasets. We used a liberal whole-brain mask with a threshold of 0.1 and a first-degree auto-regressive model to remove the autocorrelation in the signal. Regressors were convolved with the canonical hemodynamic response function (HRF) of SPM12. For each individual and each model, we created contrast maps with the beta estimates of the feedback-modulated regressors representing activity in feedback blocks (Model 1), connectivity in feedback blocks (Model 2), activity in regulation onset blocks (Model 3), and connectivity in regulation onset blocks (Model 4). For Models 1 and 2, we included the following study-specific regressors to represent covariates of no interest: self-regulation training blocks (Keller et al., 2021; Zweerings et al., 2020, 2019), blocks for assessing emotional neural responses (Keller et al., 2021), for presenting a percent sign (neutral feedback) (Keller et al., 2021; Zweerings et al., 2020, 2019), for passive viewing of a picture (Keller et al., 2021; Zweerings et al., 2020) for backward counting (Hellrung et al., 2018; Zweerings et al., 2019), and for feedback related to backward counting (Hellrung et al., 2018). The same covariates were included in Models 3 and 4, as well as the remaining regulation block period for all studies. Finally, we estimated the t-value map to test for voxelwise activation differences from zero for each individual. The PPI estimation, i.e., Models 2 and 4, was not performed for the data of Scheinost et al., because of the absence of baseline blocks.

#### 2.4.4. Group-level analysis of activation and connectivity estimates

To investigate group estimates of whole-brain activation and connectivity, we performed second-level random-effects analyses using individual SPM t-value maps. We used t-values instead of the more conventional contrast maps (which here correspond to beta activation maps) to make the datasets more comparable.

Second-level one-sample t-tests were performed separately for each model, using a constant regressor as the regressor of interest to test for non-zero voxel-wise group estimates. Binary regressors representing studies (i.e., with values of one or zero indicating inputs belonging or not belonging to a given study) were included as covariates of no interest. These covariates were orthogonalized with respect to the constant regressor. We used an explicit mask that included the entire cerebrum and the superior part of the cerebellum. As specified in the preregistration, to generate whole-brain group-level thresholded maps, we applied a voxel-level inclusion threshold of p < 0.001 and a cluster-level threshold of p < 0.05, FWE (family-wise error)-corrected for multiple comparisons using random-field theory (Worsley et al., 1996). Since many brain clusters were identified as significant in Model 1, we applied a more stringent statistical threshold (p < 0.05, FWE-corrected at the voxel level) to this model in order to pinpoint regions with the strongest associations. We generated whole-brain maps for visualization using bspmview (bobspunt.com/software/bspmview/). We reported peak coordinates (multiple peaks were reported for the same clusters if the separation between them was greater than 10 mm) and the results were automatically labeled using Automated Anatomical Labelling 3 (AAL3) (Rolls et al., 2020). For visualization purposes only and due to the lack of significant results for Models 3 and 4, we also report whole-brain maps thresholded for small effect sizes for these models. These liberal maps were obtained by thresholding absolute t-values at *t* = 2.46 for Model 3 and *t* = 2.38 for Model 4, which would correspond to the threshold for a weak effect size (*d* = 0.20), obtained by the following formula:

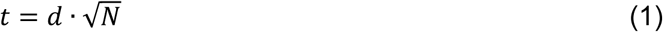

where *d* is the effect size (Cohen’s d), and N is the sample size (*N* = 152 for Model 3 and *N* = 142 for Model 4).

To establish associations between our whole-brain results and the cognitive significance of the findings, we computed the spatial Pearson’s correlation between the group-level t-value maps and 400 meta-analytic maps from Neurosynth (Yarkoni et al., 2011) using Python. This analysis was performed using the unthresholded group-level maps from Model 1, due to the hypothesis-driven positive results (Section 3.1), and Model 2, due to the exploratory-driven positive results (section 3.2) (the lack of statistical power in the PPI analysis may have contributed to the whole-brain negative results in the hypothesis-driven analysis – see Section 4.2). The 400 meta-analytic maps were obtained using the NiMARE library, which employs the Latent Dirichlet Allocation approach to the abstract or text of articles in the Neurosynth database (Poldrack et al., 2012). We then ranked and reported the 20 term sets from the meta-analytic maps with the highest positive correlations. With these rankings, we aimed to identify the main neural functional aspects related to the resulting whole-brain maps of these models.

#### 2.4.5. ROI analysis

An additional exploratory ROI analysis (i.e., not preregistered) was performed to investigate whether activation or connectivity in regions that could be potentially related to feedback processing or regulation recalibration in neurofeedback experiments, according to previous literature. We included the following ROIs involved in (1) performance-related reward processing (Camara et al., 2009; Drueke et al., 2015; Marco-Pallarés et al., 2007; Tricomi and DePasque, 2016): caudate nucleus, putamen, thalamus, NAcc, and substantia nigra; (2) self-regulation: (Sitaram et al., 2017): subgenual, rostral and superior anterior cingulate cortex (ACC), anterior and posterior insula, dorsolateral prefrontal cortex (dlPFC), posterior parietal cortex, and lateral occipital cortex; and (3) internally-oriented attention (Andrews-Hanna et al., 2014): medial prefrontal cortex (mPFC) and posterior cingulate cortex. ROI analyses were performed separately for Models 1-4. This analysis complements the whole-brain analyses because averaging the voxels within predefined ROIs might increase the signal-to-noise ratio, leading to higher statistical power. Because Models 2 and 4 considered connectivity with the NAcc and the dlPFC, respectively, these regions were removed from the ROI analyses for these Models. The ROIs and their center coordinates are described in Table 2. The full procedure for the definition of ROI masks is described in the second section of the Supp. Material.

**Table 2.**
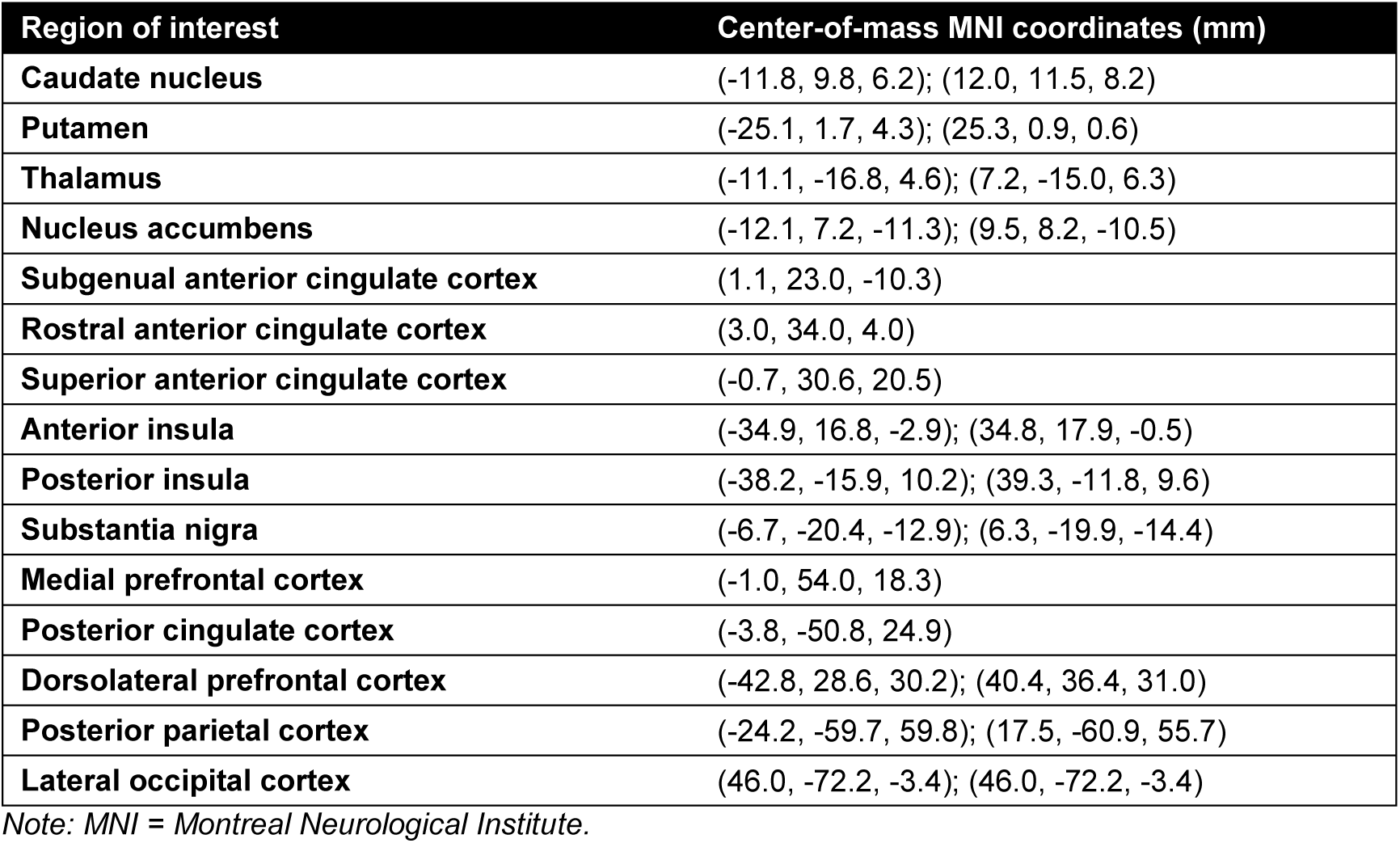
Regions of interest and their center coordinates.

Using MATLAB custom scripts, we extracted and averaged the SPM t-values, obtained in the first-level analysis, from voxels within each defined ROI. This procedure was repeated for each individual, study, and model. Due to incomplete coverage, one individual was removed from the Keller et al. dataset for the analysis of the subgenual ACC, two individuals were removed from the Hellrung et al. dataset for the analysis of the posterior cingulate gyrus; and all individuals from the Hellrung et al.’s dataset for the analysis in the posterior parietal cortex. Using R (version 4.3.2 (2022-10-31), PBC, Boston, MA, USA; rstudio.com), we ran linear regressions for each ROI and model using the following function:

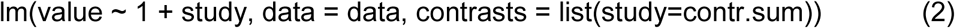

in which *value* is the mean SPM t-value within the ROI, and *study* is the study to which the individual belongs. Thus, we tested whether the intercept of the linear regression was non-null (i.e., if the association of the estimate with the feedback scores is significantly different from zero) while the studies were treated as covariates of no interest. The intercept was considered significant if the estimated 95% confidence interval of the result did not cross the zero-level. These confidence intervals were adjusted for multiple comparisons using the Bonferroni method at the ROI level. Using R, forest plots were generated for visualization.

## 3. Results

### 3.1. Model 1 – association between feedback and activation during feedback blocks

In the whole-brain analysis, we found a positive association between performance-related feedback and activity in clusters that included the basal ganglia (NAcc, ventral part of the caudate nucleus, anterior part of the putamen, and ventral pallidum), bilateral IFG, rostral ACC, PCC, mPFC, left angular gyrus, and bilateral cerebellum (voxel-wise uncorrected p < 0.001 and cluster-wise FDR-corrected p < 0.05; Fig. 1A, Table S3) during the feedback blocks. Using a more stringent threshold than the preregistered one, we found that the most robust associations were in the bilateral NAcc, mPFC, and right cerebellum (FWE-corrected voxel-level threshold p < 0.05; Fig. 1B, Table 3). In the ROI analysis, we also observed this positive association in the NAcc, rostral ACC, and mPFC (Fig. 1C). From the main meta-analytic terms associated with the result for Model 1 (Table 4), we observed that the group-level whole-brain result corresponds with brain patterns commonly linked to neurofeedback training processes, including cognitive regulation, strategic thinking, motivational systems, decision-making, retrieval processes, associative learning, and reward processing. These associations indicate that the whole-brain group-level map for Model 1 reflects not only feedback processing but also other higher-order cognitive processes involved in regulation strategies during neurofeedback training.

**Figure 1.**
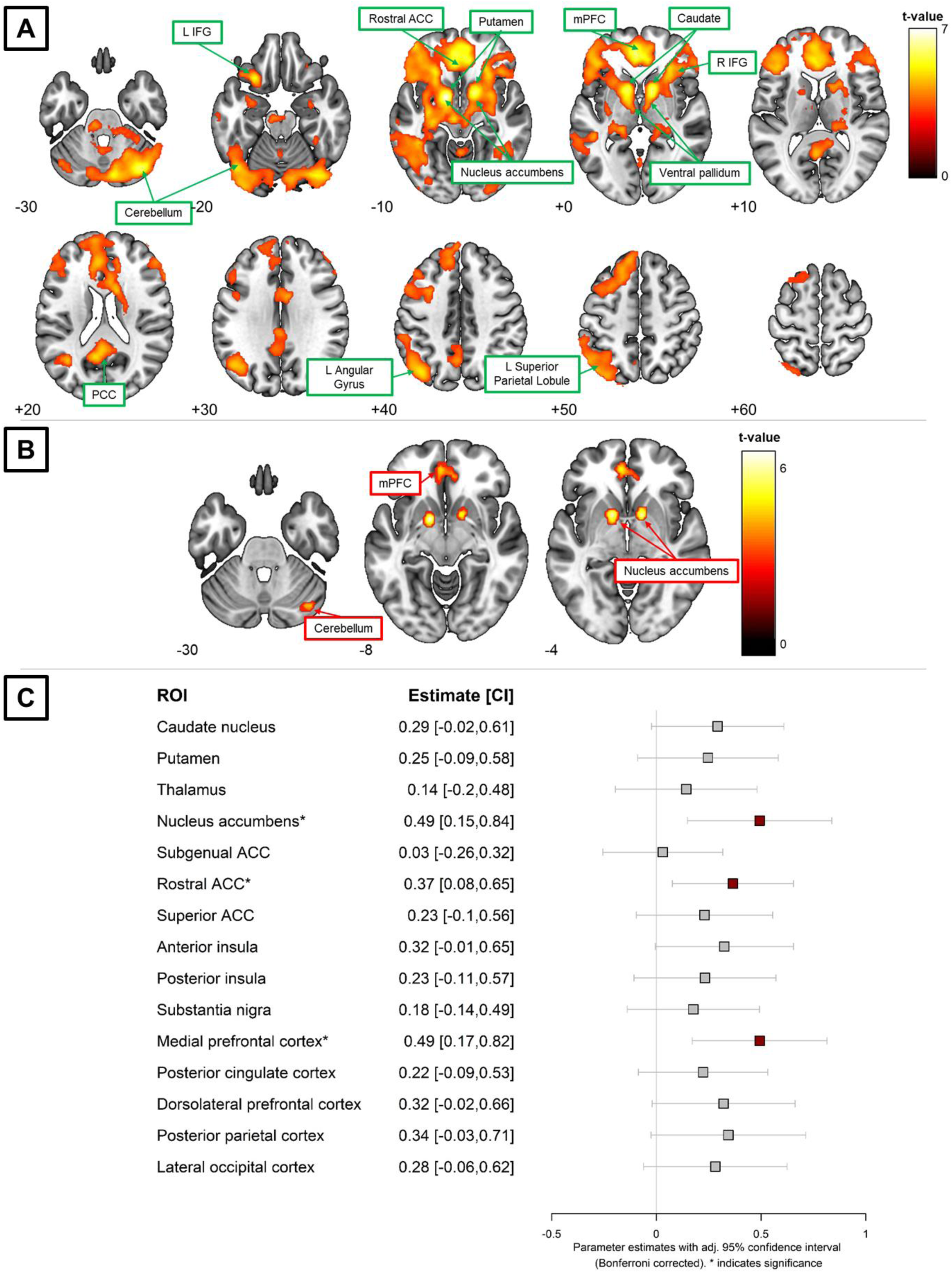
Model 1. (A) Whole-brain map (N = 153) showing brain areas where activation is positively associated with feedback scores during feedback blocks (voxel-wise uncorrected p < 0.001 and cluster-wise FDR-corrected p < 0.05). The z-coordinates for the axial slices are displayed at the bottom left corner of each slice. (B) Using a more stringent threshold than the preregistered one, we found that the most robust associations were in the nucleus accumbens, medial prefrontal cortex, and cerebellum (FWE-corrected voxel-level threshold p < 0.05). (C) Results from the region-of-interest (ROI) analysis showing regions where activation is positively associated with feedback values during feedback blocks. Asterisks and red color represent significant positive associations, while gray represents no significant association. CI = confidence interval; ACC = anterior cingulate cortex.

**Table 3.**
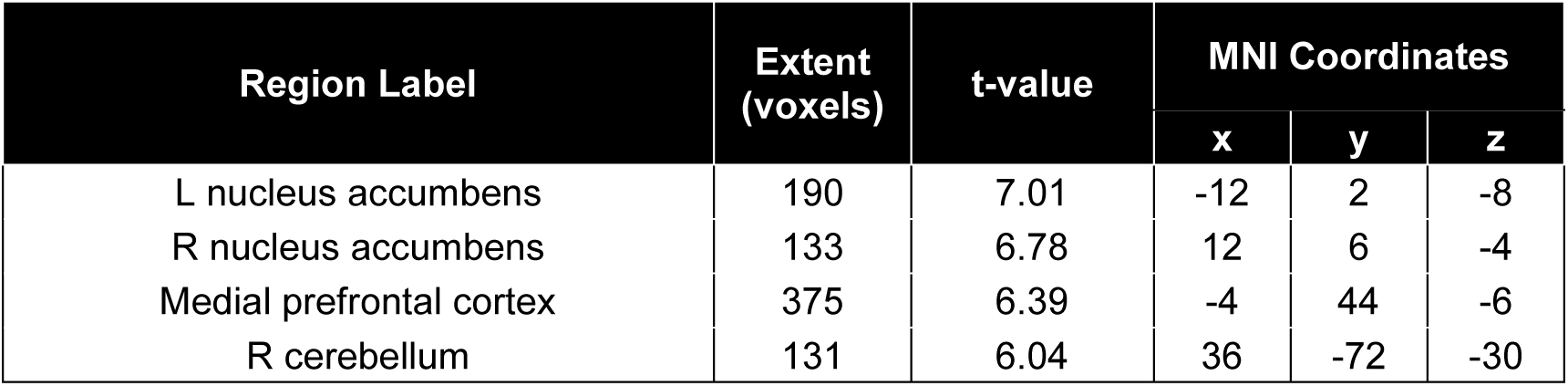
Significant clusters in whole-brain analyses for Model 1 (Fig. 1B, strong association between activity and feedback scores). Regions were labeled based on the meta-analytic associations of Neurosynth.

**Table 4.** Top meta-analytic terms for the highest correlations between group-level t-value maps (for Models 1 and 2) and meta-analytic maps. These computations are based on spatial correlations using Pearson’s correlation across 400 meta-analytic maps from Neurosynth (Yarkoni et al., 2011). Highlighted text shows meta-analytic terms that can be associated to cognitive processes typically related to neurofeedback training.

| Rank | Model 1 – association between performance-related feedback and activity |  | Model 2 – association between performance-related feedback and connectivity |  |
| --- | --- | --- | --- | --- |
|  | Correlation | Meta-analytic terms | Correlation | Meta-analytic terms |
| 1 | 0.33 | **retrieval, memory, episodic** | 0.26 | frontal, inferior, gyrus |
| 2 | 0.31 | items, recognition, item | 0.21 | prefrontal, cortex, dorsolateral |
| 3 | 0.30 | pfc, prefrontal, cortex | 0.21 | face, faces, fusiform |
| 4 | 0.30 | encoding, memory, recognition | 0.20 | **error, errors, monitoring** |
| 5 | 0.30 | prefrontal, cortex, dorsolateral | 0.19 | **prediction, predictive, predictions** |
| 6 | 0.29 | **motivation, avoidance, approach** | 0.18 | inhibition, response, stop |
| 7 | 0.29 | pairs, associative, associations | 0.18 | resonance, magnetic, control |
| 8 | 0.29 | mpfc, medial, prefrontal | 0.18 | faces, face, facial |
| 9 | 0.29 | **decision, making, choice** | 0.18 | category, categories, categorization |
| 10 | 0.27 | pictures, neutral, picture | 0.18 | **task, performing, cognitive** |
| 11 | 0.26 | **strategies, strategy, strategic** | 0.17 | reading, letter, chinese |
| 12 | 0.25 | relational, strength, strong | 0.17 | reading, phonological, dyslexia |
| 13 | 0.25 | emotional, emotion, amygdala | 0.17 | detection, novelty, oddball |
| 14 | 0.24 | deception, truth, lying | 0.17 | **conflict, response, monitoring** |
| 15 | 0.24 | **regulation, emotion, reappraisal** | 0.16 | pictures, neutral, picture |
| 16 | 0.24 | cortex, anterior, cingulate | 0.16 | cortex, visual, temporal |
| 17 | 0.24 | **learning, learned, associations** | 0.16 | memory, working, verbal |
| 18 | 0.24 | **reward, striatum, monetary** | 0.16 | deception, truth, lying |
| 19 | 0.24 | cortex, lateral, prefrontal | 0.15 | rule, rules, abstraction |
| 20 | 0.23 | rule, rules, abstraction | 0.15 | encoding, memory, recognition |

### 3.2. Model 2 – association between feedback and connectivity with NAcc during feedback blocks

In the whole-brain analysis, we found no associations between performance-related feedback and task-related connectivity with the NAcc during the feedback blocks. (For illustration purposes only, we provide a map showing associations at a lower threshold than the preregistered one; Fig. S1, Table S4). In the ROI analysis, we observed a positive association in the superior ACC, bilateral anterior insula, and substantia nigra (Fig. 2). From the main meta-analytic terms associated with the result for Model 2 (Table 4), we observed that the whole-brain group-level map corresponds with brain patterns commonly related to error and conflict monitoring, predictive processing, and cognitive performance. These associations indicate that the whole-brain group-level map for Model 2 reflects cognitive control towards task performance.

**Figure 2.**
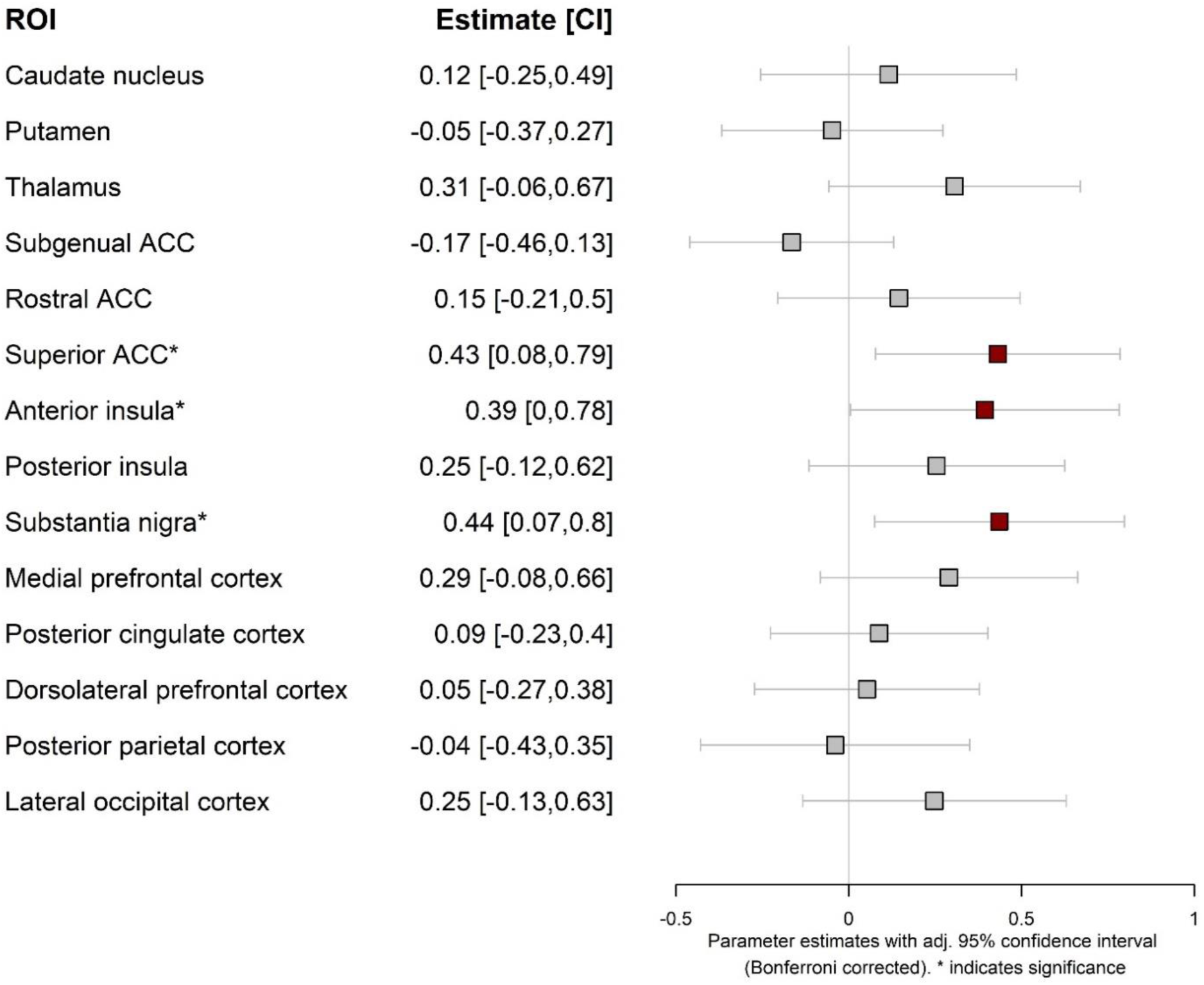
Results from region-of-interest (ROI) analysis of Model 2 showing regions whose connectivity with NAcc is positively associated with feedback values during feedback blocks. Asterisks and red color represent significant positive associations and gray color represent no significant association. CI = confidence interval, ACC = anterior cingulate cortex.

### 3.3. Model 3 – association between feedback and activation during self-regulation

We found no associations between the preceding performance-related feedback and activity during regulation blocks in the whole-brain analysis. For visualization, a whole-brain map showing effect sizes greater than small for this association is provided (Fig. S2A, Table S5). Similarly, no associations were found between the preceding level of performance-related feedback and activity during regulation blocks in the ROI analysis (Fig. S2B).

### 3.4. Model 4 – association between feedback and connectivity with the dlPFC during self-regulation

We found no associations between the preceding performance-related feedback and task-related connectivity with the dlPFC during regulation blocks in the whole-brain analysis. For visualization, a whole-brain map showing effect sizes greater than small for this association is provided (Fig. S3A, Table S6). Similarly, no associations were found between the preceding level of performance-related feedback and task-related connectivity with the dlPFC during regulation blocks in the ROI analysis (Fig. S3B).

## 4. Discussion

We investigated brain regions whose activity and connectivity were associated with feedback processing or regulation recalibration during neurofeedback training. The findings are based on a mega-analysis of eight studies and a total of 153 individuals. We found positive associations between activity and feedback scores during feedback processing in the nucleus accumbens (NAcc), putamen, caudate, ventral pallidum, medial prefrontal cortex, bilateral inferior frontal gyrus, rostral anterior cingulate cortex, posterior cingulate cortex, left angular gyrus, left superior parietal lobule, and cerebellum (Fig. 1). Additionally, an exploratory connectivity analysis revealed that connectivity between the NAcc and several brain regions – specifically the substantia nigra, anterior insula, and superior anterior cingulate cortex – was positively associated with feedback scores during feedback processing (Fig. 2). We found no regions whose activity or connectivity with the dlPFC was significantly associated with feedback scores during regulation recalibration. In this discussion, we will elaborate on our findings of brain regions associated with reward processing during neurofeedback training by grouping them into core functional brain networks, namely the basal ganglia, the default mode network, and the salience network, as well as other regions associated with attentional control and activity-modulating dopaminergic regions. A summary of the findings and possible feedback-related roles of each network is depicted in Figure 3.

**Figure 3.**
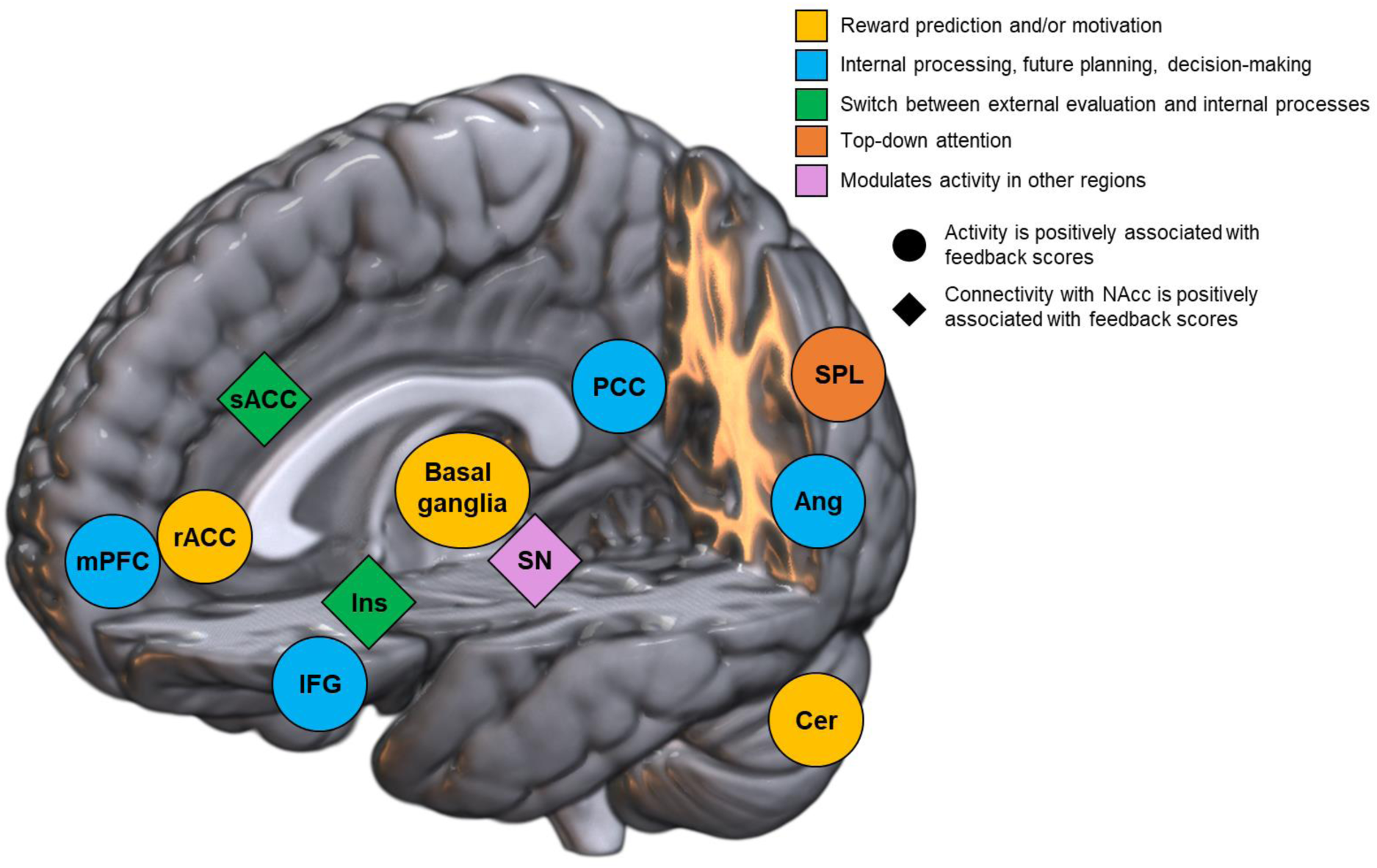
Summary of the findings and suggested interpretation in the context of neurofeedback training and feedback. The colors represent key functions associated with the corresponding brain regions and presumably employed during neurofeedback training. These functional groupings are based on established associations from previous studies, described in sections 4.1 and 4.2. The shapes represent whether the finding is related to either activity or connectivity estimations in the analysis. mPFC = medial prefrontal cortex, rACC = rostral anterior cingulate cortex, sACC = superior anterior cingulate cortex, IFG = inferior frontal gyrus, Ins = insular cortex, SN = substantia nigra, PCC = posterior cingulate cortex, Ang = angular gyrus, SPL = superior posterior lobule, Cer = cerebellum, L/R = left/right.

In neurofeedback training, individuals can learn control over, e.g., activity within a predefined region by receiving contingent feedback that reflects the regulation performance. Several theories have been postulated to explain feedback-based learning during neurofeedback training. For example, it was proposed that operant conditioning (i.e., reinforcement) drives learning in neurofeedback training (Hellrung et al., 2022; Lubianiker et al., 2022; Sitaram et al., 2017). After the feedback is processed and evaluated, the individual’s strategy is modulated in order to improve performance and prediction error is thought to decrease over successful training. For example, a “neurofeedback control network” has been proposed based on contrasting self-regulation over sham feedback (Ninaus et al., 2013), on the identification of common activations during regulation blocks across fMRI-neurofeedback studies (Emmert et al., 2016), or based on the comparison between continuous and intermittent feedback (Dewiputri et al., 2021). These studies indicate the involvement of regions comprising the salience and the frontoparietal control networks, related to the control of cognitive processes modulated by bottom-up stimuli and the switch between externalized and internalized cognitive processes (Corbetta and Shulman, 2002; Dosenbach et al., 2008; Menon, 2011; Sridharan et al., 2008). All these processes represent important aspects of neurofeedback learning, and revealing their underlying neural mechanisms is paramount to a better understanding of feedback-based learning.

### 4.1. Model 1 – association between feedback and activation during feedback blocks

We confirmed our first hypothesis, which stated that, during feedback presentation blocks, performance-related feedback scores are positively associated with activity in reward-related brain regions. More specifically, we observed that brain activity associated with feedback processing was mainly located in the bilateral NAcc, which is part of the basal ganglia and an important region of the dopaminergic reward system. The fact that the reward-related regions are associated with feedback scores provided during neurofeedback training supports the theory that neurofeedback learning is driven by reinforcement learning (Hellrung et al., 2022; Lubianiker et al., 2022; Shibata et al., 2019). We also observed this association in the medial prefrontal cortex, rostral anterior cingulate cortex, and cerebellum (Fig. 1A and 1C); as well as in other parts of the basal ganglia, the bilateral inferior frontal gyrus, posterior cingulate cortex, left angular gyrus, and left superior parietal lobule (Fig. 1A).

The basal ganglia have been linked to reward (Nieuwenhuis et al., 2005b; Schultz, 2016; Yacubian et al., 2006), learning (Camara et al., 2009; Schultz, 1998), and motivational processes (Ikemoto et al., 2015). Animal studies show that an intact basal ganglia response to reward is essential for successful learning in neurofeedback training (Koralek et al., 2012). Previous studies have suggested that the basal ganglia are involved in regulating brain activation in neurofeedback studies (Dewiputri et al., 2021; Emmert et al., 2016; Hinterberger et al., 2005; Shibata et al., 2019). However, such findings on self-regulation could potentially be entangled with feedback processing, as the analyses were performed in neurofeedback experiments employing continuous feedback (Haller et al., 2013). During neurofeedback experiments with continuous feedback, the evaluation of feedback and regulation of the target brain regions occurs simultaneously. Here, we analyzed data from neurofeedback experiments that temporally segregate feedback appraisal and regulation phases. Therefore, we suggest that basal ganglia involvement is related to feedback appraisal, rather than self-regulation of neurofeedback training. We observed a strong association between feedback scores and activity during feedback blocks in the NAcc (part of the ventral striatum). This association is consistent with several studies showing that the NAcc shows higher activity for positive compared to negative feedback (Delgado et al., 2000; Fouragnan et al., 2018; Marco-Pallarés et al., 2007; Nieuwenhuis et al., 2005b; Yacubian et al., 2006). The NAcc is an integrative region that connects cortical and limbic regions, as well as the midbrain, and regions associated with dopamine release (Garris et al., 1999). Other studies have shown that activity in the NAcc correlates with gain-related prediction error (Montague et al., 1996; Shohamy, 2011; Yacubian et al., 2006), which is crucial for associative learning (Daniel and Pollmann, 2012). The involvement of the NAcc in feedback processing during neurofeedback, and its established link to prediction error may also indicate the applicability of the associative learning theory of neurofeedback learning. The ventral striatum has also been implicated in unconscious reward processing (Ramot et al., 2016; Sitaram et al., 2017). Therefore, it is noteworthy that neurofeedback studies using implicit feedback, i.e., a setup that operates without the conscious awareness of the participant but with indirect methods to induce intended brain activity (Watanabe et al., 2017), also elicit activation of the ventral striatum (Shibata et al., 2019).

Although the highest association between activity and feedback scores during neurofeedback training was found in the NAcc, we also found this association in other regions of the basal ganglia (Fig. 1A). We observed this positive association in the ventral part of the caudate nucleus and the anterior part of the putamen. These regions comprise the dorsal striatum, which has been reported to be more activated in response to positive than negative feedback (Drueke et al., 2015; Duijvenvoorde et al., 2008; Wächter et al., 2009). It has been suggested that the dorsal striatum receives reward-related information from the ventral striatum and uses this information to predict and maximize positive outcomes (Tricomi and DePasque, 2016; Yin et al., 2005). The dorsal striatum is part of the “associative loop”, which links rewards or punishments with previous actions (Tricomi and DePasque, 2016). Furthermore, activation in the caudate due to performance-related intrinsic feedback is similar to that produced by extrinsic (e.g., monetary) rewards or punishments (Tricomi and DePasque, 2016). Tricomi et al. (2006) proposed that the caudate facilitates feedback-based learning by identifying and assigning value to correct and incorrect responses (Tricomi et al., 2006). The ventral caudate has also been associated with short-term reward and behavioral learning through prediction error (Haruno et al., 2004). Furthermore, the stimulus-action-reward association was found to be located in the anterior part of the putamen (Haruno and Kawato, 2006). These fMRI findings regarding a more precise anatomical definition of the ventral striatum in reward processing are consistent with our results. We also observed that the anterior part of the ventral pallidum (Fig. 1A), another major basal ganglia region, was positively associated with feedback scores during neurofeedback training. Studies indicate that the ventral pallidum codes several aspects of reward, such as information about prediction error, valence, and surprise (Schultz, 2016; Tachibana and Hikosaka, 2012). Furthermore, we observed a positive association between feedback scores and activity in the rostral anterior cingulate cortex. The present findings are consistent with previous research indicating that the rostral anterior cingulate cortex is influenced by a positive discrepancy between actual and anticipated feedback, and that this region plays a pivotal role in evaluating salient feedback and shaping optimal learning strategies (Amiez et al., 2012).

Our meta-analytic association analysis suggests that the results of Model 1 capture not only feedback processing, but also other higher-order cognitive processes involved in neurofeedback training (Table 4). Highly ranked meta-analytic terms – that can be summarized broadly as cognitive regulation, strategic thinking, motivational systems, decision-making, retrieval processes, and associative learning – indicate that the resulting whole-brain map may also relate to the process of identifying and optimizing mental states or strategies that yield high feedback scores. These processes are typically associated with cortical rather than subcortical activations. Indeed, we observed a positive association between activity and feedback scores during neurofeedback training in key regions of the default mode network (Andrews-Hanna et al., 2014): mPFC, posterior cingulate cortex, and left angular gyrus (Fig. 1A). This large-scale brain system, which is commonly associated with deactivation in response to stimuli demanding externally-directed attention (Fox et al., 2005), conversely shows activation when internally-directed attention is induced (Gusnard et al., 2001; Harrison et al., 2008; McDonald et al., 2017; Pamplona et al., 2020; Spreng, 2012). Previous findings support that the presentation of feedback during neurofeedback training induces the involvement of some regions of the default mode network (Shibata et al., 2019) or even the whole network (Radua et al., 2018). Interestingly, our results show that the more positive the performance-related feedback, the greater the activity in the default mode network. This finding indicates that positive rewards resulting from successful self-regulation during training may elicit higher levels of internally-focused attention, potentially related to the evaluative aspects of the strategy employed. Alternatively, or possibly in conjunction with this, negative feedback may prompt an active, goal-directed control process that alters the self-regulation strategy, drawing on higher-order cognitive processes and suppressing the default mode network. The process of revisiting, evaluating, and selecting self-regulatory strategies essentially uses internally-focused attention (Kam and Handy, 2013). An internal evaluation of the self-regulatory strategy is presumably necessary to maximize performance in subsequent trials and thus optimize reward. The association between default mode network activity and feedback scores may also support the reinforcement learning theory in neurofeedback: higher rewards elicit a higher level of internal appraisal, which in turn may be beneficial to drive learning over training. Methodologically, our findings also suggest that caution should be taken when designing neurofeedback experiments that target voluntary deactivation of the default mode network (or the other regions resulting from Model 1) using continuous feedback. While down-regulation of this brain system is intended, positive feedback generated by successful regulation could, at least partially, elicit positive responses and thus operate in the opposite direction as intended. Therefore, this feedback presentation interference favors the use of intermittent over continuous feedback, depending on the target region.

Associations in other cortical regions evidencing higher-order cognitive processes during neurofeedback training were observed. Our results show that activity in cortical regions, such as the bilateral inferior frontal gyrus, was positively associated with feedback scores during neurofeedback training (Fig. 1A). These findings are consistent with previous studies on the neural mechanisms of reward (Duijvenvoorde et al., 2008; Radua et al., 2018) and strategy execution during intermittent neurofeedback (Dewiputri et al., 2021). Involvement of the inferior frontal cortex may reflect the semantic conceptualization of abstract (nonverbal) information (Hoffman et al., 2015) or the selection of competing executed strategies retrieved from semantic memory during feedback presentation (Thompson-Schill et al., 1997). Reward-related activation in the visual cortex has been previously reported and interpreted as enhanced visual processing of stimuli (Drueke et al., 2015). We also observed that the cerebellum was positively associated with feedback scores. While relatively little is known about cerebellar activations and feedback processing, activation in the right cerebellum has previously been associated with positive (compared to negative) feedback (Marco-Pallarés et al., 2007). A recent animal study also showed that the cerebellum sends excitatory projections to reward-encoding regions (Carta et al., 2019). Finally, we found that feedback scores were positively associated with activity in the left superior parietal lobule (Fig. 1A). This region has been implicated in top-down attentional control processes (Corbetta and Shulman, 2002; Green and McDonald, 2008), goal-directed behavior (Bressler and Menon, 2010), and feedback processing (Crone et al., 2008). Altogether, we interpret this region to be responsible for controlling attentional effort in proportion to feedback values.

### 4.2. Model 2 – association between feedback values and connectivity with NAcc during feedback blocks

Model 1 confirmed the NAcc as the key region for feedback processing during neurofeedback training. Model 2 investigated regions whose connectivity with the NAcc was modulated by the feedback scores. This was done using PPI, which controls for functional connectivity that is independent of the task (e.g., in resting-state designs), thus revealing only functional connectivity that is modulated by feedback processing (O’Reilly et al., 2012). We confirmed our second hypothesis, which stated that, during feedback presentation blocks, performance-related feedback scores are positively associated with connectivity within reward-related brain regions. More specifically, significant associations were identified between feedback scores and connectivity with the NAcc in the substantia nigra, anterior insula, and superior anterior cingulate cortex (Fig. 2). It is important to note that this confirmation is based on exploratory analysis, specifically ROI analysis, rather than whole-brain analysis. However, a corollary of the definition of PPI analysis is that it lacks power and is prone to a high rate of false negatives (O’Reilly et al., 2012), primarily due to the similarity between the PPI regressor and both the task and seed time-courses. (In fact, by applying a more liberal threshold, it is possible to partially reproduce the regions identified in the ROI analysis within the whole-brain map; Fig. S1).

It is known that the substantia nigra connects to the NAcc via the dopaminergic pathway and projects to various sites of the basal ganglia (Camara et al., 2009; Rabey and Hefti, 1990; Schultz, 2016). In fact, reward processing has been associated with activation in the substantia nigra (Cohen et al., 2012; Yasuda et al., 2012). The anterior insula and the superior anterior cingulate cortex (Fig. 2) constitute the salience network, which facilitates switching between internally- and externally-oriented cognitive processes (Menon, 2011; Sridharan et al., 2008). In this regard, our meta-analytic association analysis implicates that the group-level whole-brain map of Model 2 relates to monitoring of prediction error towards task performance. These findings align with a recent study reporting that the salience network underlies feedback processing during intermittent neurofeedback (Dewiputri et al., 2021). In the context of feedback-related reward, the recruitment of the salience network may be necessary to mediate the external reward evaluation (i.e., prediction error monitoring), followed by the internal weighting and judgment of adopted mental strategies to optimize feedback (i.e., task performance). The functional coupling between the NAcc and the salience network indicates that the switching of mental processes undertaken by the salience network may be triggered by basal ganglia activity and the positive association indicates that this triggering is stronger for positive feedback.

### 4.3. Models 3 and 4 – null results for associations between feedback values and activation/dlPFC connectivity during self-regulation

Models 3 and 4 were designed to investigate activation/connectivity during the regulation blocks associated with the preceding feedback score. Our aim was to investigate how feedback can modulate regulation recalibration, specifically strategy adaptation during self-regulation after receiving feedback. We hypothesized that there would be negative associations with feedback due to its increased adaptation-demanding nature of negative feedback. We hypothesized that this association would be primarily located in the dlPFC. The dlPFC has been suggested to be involved in cognitive control during neurofeedback learning (Ninaus et al., 2013; Sitaram et al., 2017). However, we could not confirm our third and fourth hypotheses, which stated that, during regulation blocks, performance-related feedback scores would be associated with activity/connectivity with dlPFC in brain regions related to neurofeedback control. Therefore, although activation in the dlPFC during self-regulation has been found to be ubiquitous across studies (Emmert et al., 2016), its initial-phase activation or functional coupling with other regions may not be modulated by feedback scores. Another possibility is that the process we hypothesized to take place at the beginning of the regulation blocks may have already been initiated when the performance-related feedback was presented. This interpretation would explain why we did not observe time-locked responses in the subsequent regulation blocks. In this case, recalibration could have already been occurring during the feedback blocks, and the results in Models 1 and 2 would reflect not only feedback processing but also adaptation or reinforcement of mental strategies. In fact, the results from the meta-analytic association analysis (Table 4) suggest that this was in fact the case: results from Models 1 and 2 were not restricted to feedback processing but also higher-order cognitive processes related to the identification and refinement of regulation strategies during neurofeedback training.

The regulation recalibration in our study relies on the implicit assumption that participants could both learn the regulation task and recalibrate regulation effectively during early training phases, since all feedback blocks and subjects were included in the analysis. Ideally, a more precise model would focus exclusively on successful runs (or even blocks) or from participants identified as learners. However, defining “learners” or “successful runs” is challenging, as the definition of learning success lacks consensus in the literature (Haugg et al., 2020). From an analytical perspective, we used available feedback scores across all training, since restricting the analysis to successful blocks alone would significantly reduce the amount of data, potentially leading to insufficient data for a regression analysis. This generalization may partly explain our negative findings for Models 3 and 4, as the inclusion of blocks where regulation based on feedback was not achieved, as well as the inclusion of non-learners, may have obscured positive results. In addition, factors like noisy feedback signals and low individual learning rates may have further hindered the findings. However, our primary focus was to characterize prediction error, where discrepancies between expected and actual feedback drive learning. This concept remains consistent throughout the learning process, independent of phase or individual success. Future research could refine this approach by evaluating feedback-dependent regulation recalibration in only successful blocks and learners.

### 4.4. Limitations of the study

Firstly, the feedback-modulated connectivity analyses were based on one seed each for Models 2 and 4. Due to our pre-registered methodology and computationally demanding analysis, we restricted our analysis to a priori defined seeds. Future work may investigate other potentially representative regions.

Secondly, it should be noted that feedback appraisal is inherently subjective (Tricomi and DePasque, 2016). For example, highly motivated or initially successful individuals might perceive a half-full feedback representation as more negative than those who are poorly motivated or initially unsuccessful. Similarly, in bidirectional neurofeedback studies, the ability to regulate in one direction may be easier than in the other. Therefore, an individual’s perception of success may depend on the difficulty of regulating in each direction. To account for such differences, future studies may include data on subject-specific motivation levels in their analysis.

Thirdly, only one out of eight studies included sham feedback; therefore, we could not compare sham and veridical feedback and could not extend our results to sham feedback. However, we argue that the neural mechanisms for processing sham feedback should lead to similar results, as long as the feedback presentation is not perceived as sham feedback (Ninaus et al., 2013).

Fourthly, feedback success and failure were evaluated using a one-dimensional scale and cannot be dissociated in our study. From a methodological perspective, it would not be reasonable to separate failure and success for some datasets of our study (Radua et al., 2018), since the feedback representations varied parametrically on the same scale. However, it is conceivable that some of the anticipated regions (e.g., in ROI analyses) were not identified as associated with feedback due to the modeling strategy of defining success and failure within a single one-dimensional scale. We argue that studying the neural mechanisms of graded feedback, as opposed to binary, is, therefore more informative and ecologically valid (Radua et al., 2018).

## 5. Conclusion

Our mega-analysis using data from eight fMRI-neurofeedback experiments revealed that feedback processing is primarily associated with activity in subcortical regions (NAcc, putamen, caudate, and pallidum) and the cerebellum. Such findings indicate that neurofeedback is processed in core regions of the reward system, suggesting that inherent motivational and reward/punishment aspects shape neurofeedback learning. The evoked neural responses are similar to those elicited by extrinsic or primary rewards, representing the dopaminergic release for rewarding feedback. We also observed that activity and connectivity with the NAcc was positively associated with feedback scores in several large-scale networks. This involvement is likely to represent the internally-directed attention to the strategies adopted and their appraisal for the subsequent trials, the switch between attention to internal appraisal and external reward modulated by the basal ganglia, and the top-down attention for associating feedback and self-regulatory performance. As a corollary to the positive association between activity and feedback scores, our findings have implications for neurofeedback paradigms that use continuous feedback to train down-regulation of the brain regions reported here. Specifically, feedback reward associated with successful down-regulation would elicit a positive neural response to down- regulation effort, i.e., researchers may consider a possible interference between down-regulation performance and feedback-related activation.

Our findings contribute to the understanding of how self-regulatory learning is promoted by neurofeedback paradigms. We provide evidence that this learning process occurs through reinforcement learning: positive feedback elicits activity in the reward system, which in turn promotes performance improvement over training. These findings may extend to other feedback-dependent learning paradigms and recalibration of strategies towards a goal (Tricomi and DePasque, 2016). In addition, we show that large-scale networks, which allocate and modulate attentional resources to both externally-presented feedback and evaluative processing of self-regulatory strategies, are involved in the learning process of neurofeedback training. Such a finding may be more specific to neurofeedback paradigms due to their introspective nature of evaluating internal self-regulatory strategies. Overall, our findings highlight the importance of feedback as a driving force for learning – from grades on a school exam to complex experimental paradigms such as neurofeedback.

## Acknowledgements

We thank Dr Samantha Fede for providing information about neurofeedback studies using intermittent feedback and Dr Bruno H. Vieira for statistical support.

## Ethics Approval Statement

This study was approved by the University of Vienna (EK 00621).

## Data and Code Availability Statement

All obtained results and scripts used for the data analysis are available on the public GitHub repository: https://github.com/gustavopamplona/FeedbackReward.

## Author Contributions

GSPP: conceptualization, methodology, software, formal analysis, investigation, data curation, writing – original draft, writing – review and editing, visualization, supervision, project administration; JZ: investigation, data curation, writing – review and editing; CL: writing – review and editing; LdE: formal analysis; ER: formal analysis, resources; AT: investigation, data curation, writing – review and editing; LH: investigation, data curation, resources, writing – review and editing; KA: investigation, data curation, writing – review and editing; DS: investigation, data curation, writing – review and editing; FK: investigation, data curation, writing – review and editing; MDR: investigation, data curation, writing – review and editing; SI: funding acquisition; SB: funding acquisition, writing – review and editing; EJH: investigation, data curation, writing – review and editing; KM: conceptualization, methodology, resources, writing – review and editing, project administration, funding acquisition; FS: conceptualization, methodology, resources, writing – review and editing, project administration, funding acquisition;

