## Supplementary material for "Neural Mechanisms of Feedback Processing and Regulation Recalibration during Neurofeedback Training"

#### List of Contents


1. Systematic search for relevant studies
2. Procedure for the definition of ROI masks
3. Figures
   - 3.1. Figure S1
   - 3.2. Figure S2
   - 3.3. Figure S3
4. Tables
   - 4.1. Table S1
   - 4.2. Table S2
   - 4.3. Table S3
   - 4.4. Table S4
   - 4.5. Table S5
   - 4.6. Table S6

#### 1. Systematic search for relevant studies

Using the Scopus systematic search (www.scopus.com/) on 05/JAN/2022, we searched for all articles including in their full text the words "neurofeedback", "fMRI", and "intermittent", i.e., we used the following string: (ALL (fmri) AND ALL (neurofeedback) AND ALL (intermittent)) AND (LIMIT-TO (DOCTYPE, "ar")). We then filtered studies based on the following exclusion criteria:

- studies whose acquisition did not include sub-cortical areas or that covered only a few slices of the brain;
- studies containing exclusively data from individuals with a psychiatric or neurological disorder, addicted to or craving a particular substance, or suffering from phobias;
- from elderly individuals;
- studies employing meta-analysis, mega-analysis, or secondary analysis from existing data;
- introduction of software for real-time fMRI, not including the acquisition of new data;
- published more than ten years prior to the systematic search (i.e., studies dated from 2012 or earlier);
- involving the analysis of artificially generated datasets;
- concerning methodological aspects of fMRI-neurofeedback;
- involving the acquisition of data from children or adolescents;
- involving techniques other than fMRI (such as EEG, TMS, or fNIRS);
- review studies;
- not related to fMRI-neurofeedback;
- studies whose feedback could not be converted into parametric values (i.e., categorical feedback);
- fMRI-neurofeedback studies using continuous feedback.

In addition, we contacted the authors of a systematic review (Fede et al., 2020) that reported 17 fMRI-neurofeedback studies using intermittent feedback and we asked for a list of these studies (nine of them had been already found in the systematic search).

#### 2. Procedure for the definition of ROI masks

ROIs were defined based on the meta-analytic maps automatically generated in Neurosynth (Yarkoni, Poldrack, Nichols, Van Essen, & Wager, 2011; neurosynth.org) on 25/OCT/2022. In this way, the ROI definition was independent of the whole-brain results (Kriegeskorte et al., 2009). First, we downloaded the meta-analytic maps generated with the terms: “caudate”, “putamen”, “thalamus”, “nucleus accumbens”, “subgenual”, “rostral anterior”, “anterior cingulate”, “anterior insula”, “posterior insula”, “substantia”, “medial prefrontal”, “posterior cingulate”, “dlpfc”, “posterior parietal”, “lateral occipital”. The voxel size for the downloaded meta-analytic maps was 2 x 2 x 2 $mm^3$. We then spatially smoothed the meta-analytic maps with a Gaussian kernel of 6-$mm^3$ FWHM. Next, we identified the 50 voxels with the highest z-values in each hemisphere and combined them to create ROIs with bilateral clusters (i.e., each ROI contained 100 voxels, 50 in each hemisphere). Medially located regions (ACC, medial prefrontal cortex, and posterior cingulate cortex) were represented by 100-voxel single clusters, i.e., without hemispheric subdivision. Visual inspection indicated unfragmented clusters in the expected locations for all ROIs except for the posterior parietal cortex. The procedure for this region led to a second, more inferior cluster in the left hemisphere. Therefore, we repeated the procedure for this region limiting the identification of peak voxels to the superior part of the brain only.

### 3. Figures

#### 3.1. Figure S1

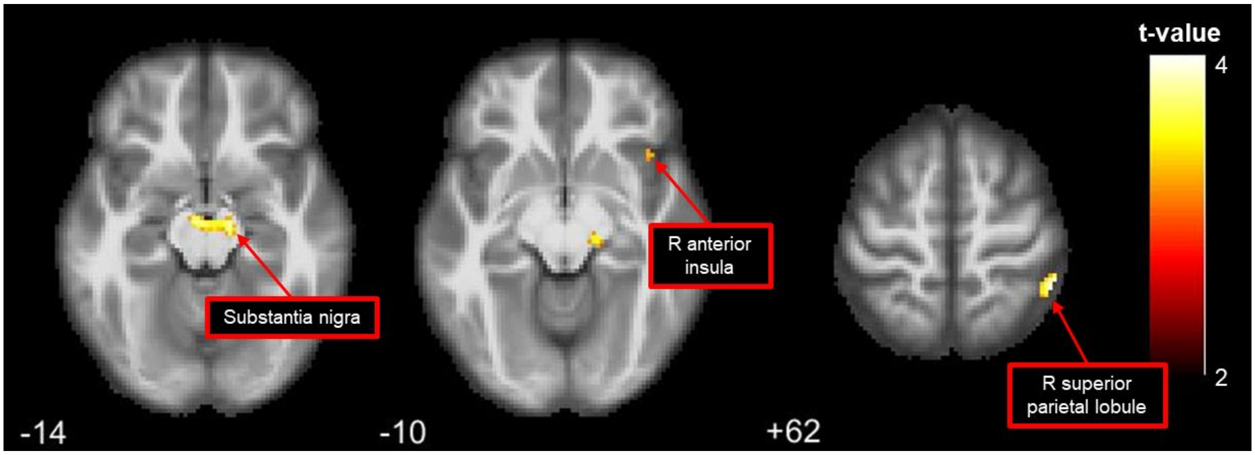


*Figure S1. For Model 2 and a lower threshold (uncorrected voxel-level threshold of p < 0.001), a whole-brain map shows brain areas whose connectivity with NAcc is positively associated with feedback scores during feedback blocks. The z-coordinates for axial slices are displayed at the left bottom corner of each slice.*

#### 3.2. Figure S2

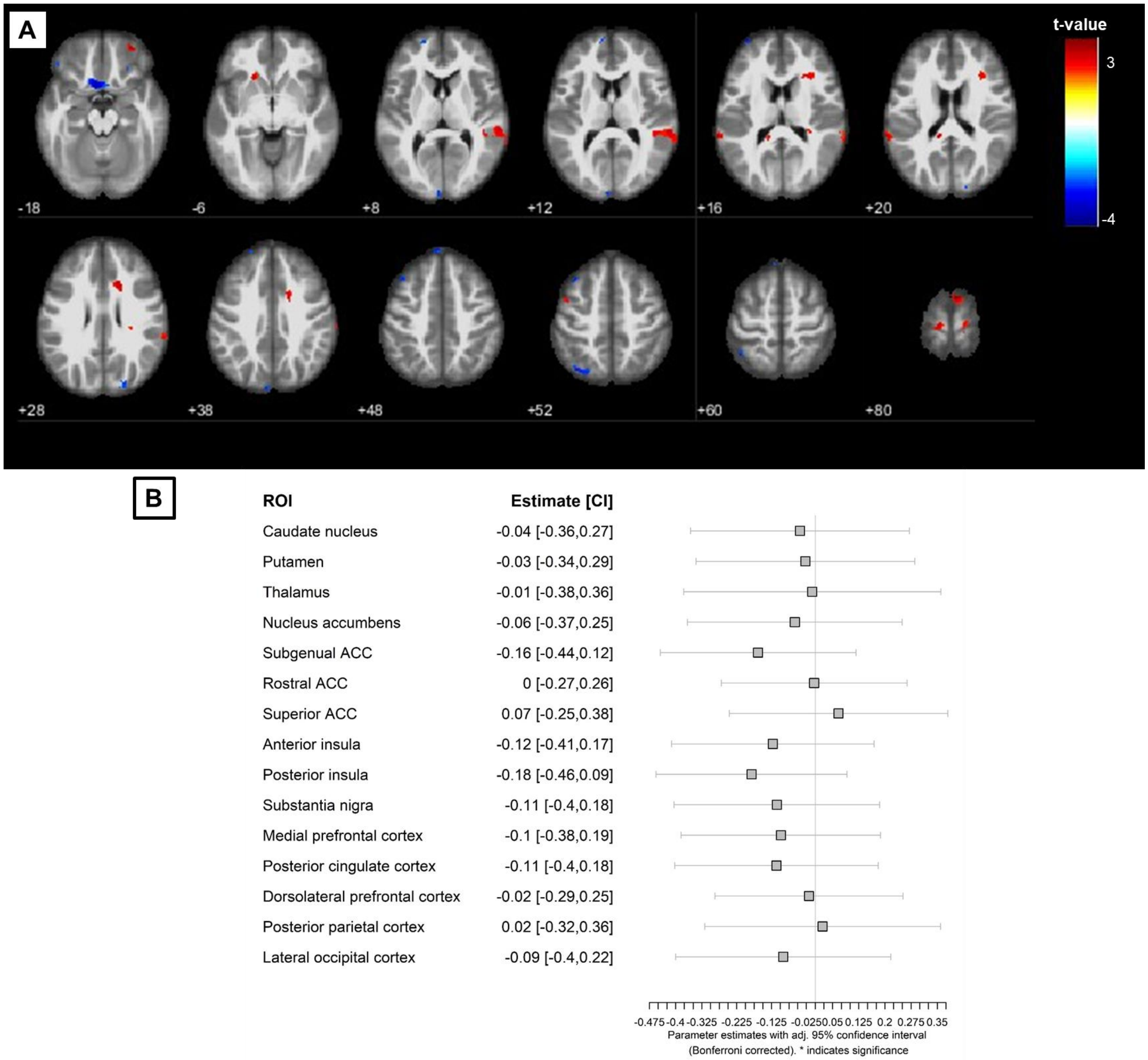


*Figure S2. Model 3. (A) Whole-brain map showing brain areas where the effect size of activation is positively (red)/negatively (blue) associated with feedback scores during self-regulation. Only voxels with absolute t-values were higher than 2.46 are displayed to not show regions where effect size was smaller than 0.2. The z-coordinates for axial slices are shown in the lower left corner of each slice. (B) Region-of-interest (ROI) analysis showed no significant association between activation and feedback values during self-regulation. CI = confidence interval, ACC = anterior cingulate cortex.*

#### 3.3. Figure S3

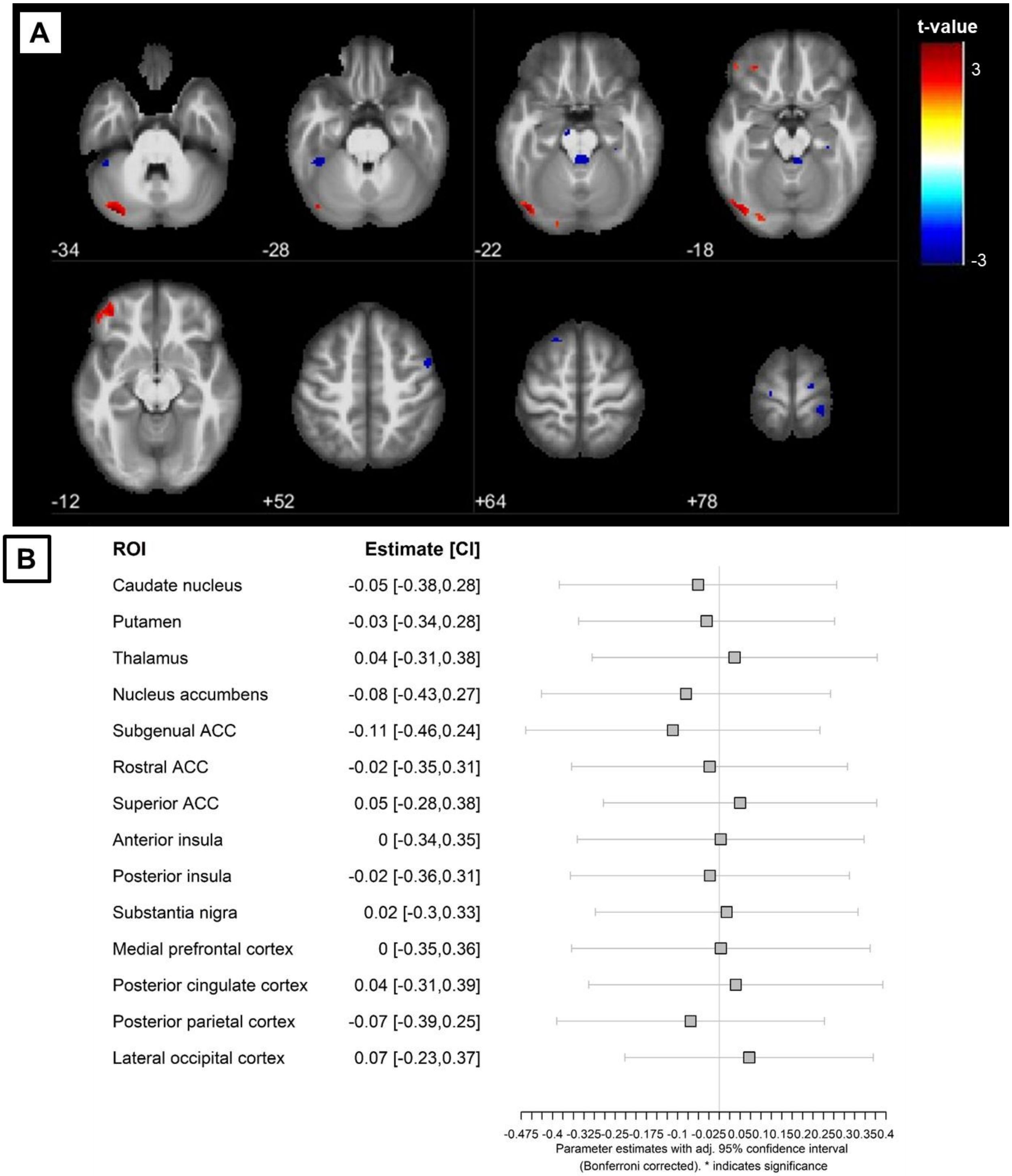


*Figure S3. Model 4. (A) Whole-brain map showing brain areas where the effect size of connectivity with the dlPFC is positively (red)/negatively (blue) associated with feedback scores during self-regulation. Only voxels with absolute t-values were higher than 2.38 are displayed to not show regions where effect size was smaller than 0.2. The z-coordinates for axial slices are shown*

*in the lower left corner of each slice. (B) Region-of-interest (ROI) analysis showed no significant association between connectivity with the dlPFC and feedback values during self-regulation. CI = confidence interval, ACC = anterior cingulate cortex.*

### 4. Tables

#### 4.1. Table S1

*Table S1. Summary of the fMRI acquisition parameters for each included study.*

| Study | Brand/ Model | Number of channels in head coil | TR (s) | TE (ms) | Matrix size (voxels) | Flip angle (°) | Number of slices | Voxel size (mm³) | Slice gap (mm) | Slice order | Slicing direction | Number of volumes |
|---|---|---|---|---|---|---|---|---|---|---|---|---|
| **Keller et al., 2021** | Siemens MAGNETOM Trio | 20 | 2 | 28 | 64 x 64 | 77 | 34 | 3 x 3 x 3 | 0.75 | Interleaved | Ascending | 230 |
| **Zweerings et al., 2019** | Siemens MAGNETOM Trio | 20 | 2 | 28 | 64 x 64 | 77 | 34 | 3 x 3 x 3 | 0.75 | Interleaved | Ascending | 230 |
| **Zweerings et al., 2020** | Siemens MAGNETOM Prisma | 20 | 2 | 27 | 64 x 64 | 90 | 34 | 3 x 3 x 3 | 0.8 | Interleaved | Ascending | 230 |
| **Hellrung et al., 2018** | Siemens MAGNETOM Trio | 12 | 2 | 25 | 64 x 64 | 90 | 32 | 3 x 3 x 2.6 | 0.26 | Interleaved | Ascending | 260 |
| **Pamplona et al., 2020** | Philips Achieva | 32 | 2 | 30 | 80 x 80 | 80 | 37 | 3 x 3 x 4 | 0.5 | Continuous | Ascending | 190 |
| **Amano et al., 2016** | Siemens Verio | 12 | 2 | 26 | 64 x 64 | 80 | 33 | 3 x 3 x 3.5 | 0 | Interleaved | Yes | 165 |
| **Scheinost et al., 2020** | Siemens Trio TIM | 32 | 2 | 25 | 64 x 64 | 90 | 34 | 3.5 x 3.5 x 4 | 0 | Interleaved | Ascending starting with 2 | 428 |
| **Krause et al., 2021** | Siemens Skyra | 32 | 1 | 33 | 88 x 88 | 60 | 64 | 2.4 x 2.4 x 2.4 | 0 | Interleaved | Ascending starting with 2 | 600 |

*Note:* TR/TE = time of repetition/ time of echo.

#### 4.2. Table S2

*Table S2. Summary of experimental design for all studies*

| Study | Experiment aspect | Description |
|---|---|---|
| **Keller et al., 2021** | General instruction for regulation blocks | "Suggested strategies were to think (1) the situation will change in the future, (2) the situation is not as bad as it looks" |
| | General instruction for baseline | None |
| | Number of training days | 2 |
| | Number of training runs | 4 |
| | Duration of the training run | Around 7 min |
| | Number of baseline blocks | 9 |
| | Duration of baseline block | 6 s |
| | Number of regulation blocks | 9 |
| | Duration of regulation block | 12 s |
| | Another condition during training run | Passively view a picture |
| | Feedback representation | 2-digit number (1 to 99) |
| | Duration of feedback block | 4 s |
| | Computation of feedback value | -- |
| | Target ROI | Left or right ventrolateral prefrontal cortex |
| | Crossover design? | Yes, with respect to the ROIs |
| | Up/downregulation | Only upregulation |
| | Stimulus during regulation block | View picture |
| | Stimulus during baseline block | Fixation cross or X |
| **Zweerings et al., 2019** | General instruction for regulation blocks | "Strategies for regulation from three categories were suggested: (1) recalling positive autobiographic memories, (2) imagining spoken language or (3) imagining a relaxing situation" |
| | General instruction for baseline | Participants had to count backward from 100 |
| | Number of training days | 2 |
| | Number of training runs | 4 |
| | Duration of the training run | 8 min |

| | | |
|---|---|---|
| | Number of baseline blocks | 9 |
| | Duration of baseline block | 6 s |
| | Number of regulation blocks | 9 |
| | Duration of regulation block | 12 s |
| | Another condition during training run | Counting |
| | Feedback representation | 2-digit number (1 to 99) |
| | Duration of feedback block | 4 s |
| | Computation of feedback value | "The BOLD percentage signal change was averaged across the two ROIs, multiplied by 100 and fed back as a positive number in the range from 1 to 99 reflecting 0–1% BOLD signal change in the up-regulation and 0 to ———1% BOLD signal change in the down-regulation condition" |
| | Target ROI | Language-related areas (left IFG and left pSTG) |
| | Crossover design? | Yes, upregulation on one day and downregulation on the other day |
| | Up/downregulation | Yes, in crossover design |
| | Stimulus during regulation block | Picture of a brain |
| | Stimulus during baseline block | Picture of a cloud |
| **Zweerings et al., 2020** | General instruction for regulation blocks | "To reduce the negative affect elicited by the aversive scenes by conceptualizing the depicted scenario " |
| | General instruction for baseline | "To respond naturally to the negative scenes" |
| | Number of training days | 1 |
| | Number of training runs | 2 (with feedback) |
| | Duration of the training run | Around 7.7 min |
| | Number of baseline blocks | 9 |
| | Duration of baseline block | 6 s |
| | Number of regulation blocks | 9 |
| | Duration of regulation block | 12 s |
| | Another condition during training run | Passively view a picture |
| | Feedback representation | 2-digit number (1 to 99) |
| | Duration of feedback block | 4 s |
| | Computation of feedback value | Reflected 0 to 1 percent signal change in the designated ROI: ([meanBOLD(reappraise) - meanBOLD(view)]× 100) |
| | Target ROI | Left lateral prefrontal cortex (lPFC) |
| | Crossover design? | Yes, two experimental (NF) and two control (NoNF) runs (feedback values correspond to only NF runs) |
| | Up/downregulation | Upregulation |
| | Stimulus during regulation block | Negative scenes |
| | Stimulus during baseline block | Negative scenes |
| **Hellrung et al., 2018** | General instruction for regulation blocks | "To perform mental strategies, such as reminiscing about personal experiences of happy situations, being with friends and sexual memories to generate positive feelings" |
| | General instruction for baseline | "Not to think about anything specific" |
| | Number of training days | 1 |
| | Number of training runs | 3 |
| | Duration of the training run | 8min40s |
| | Number of baseline blocks | 5 |
| | Duration of baseline block | 40 s |
| | Number of regulation blocks | 4 |
| | Duration of regulation block | 40 s |
| | Another condition during training run | Count backwards. Feedback also given after this condition |
| | Feedback representation | Thermometer |
| | Duration of feedback block | 4 s |
| | Computation of feedback value | Average BOLD signal from the whole previous block |
| | Target ROI | Bilateral amygdala |
| | Crossover design? | No |
| | Up/downregulation | Upregulation for happy blocks and downregulation for count blocks |
| | Stimulus during regulation block | Empty thermometer with a red up-arrow |
| | Stimulus during baseline block | Empty thermometer with an X letter |
| **Pamplona et al., 2020** | General instruction for regulation blocks | To regulate their brain activity and increase the thermometer, but attention-related instructions were given |
| | General instruction for baseline | To relax and let the mind wander |
| | Number of training days | 2 |
| | Number of training runs | 5 |
| | Duration of the training run | 6min20s |
| | Number of baseline blocks | 5 |
| | Duration of baseline block | 30 s |
| | Number of regulation blocks | 5 |
| | Duration of regulation block | 40 s |
| | Another condition during training run | No |
| | Feedback representation | Thermometer graded from blue to red |
| | Duration of feedback block | 4 s |
| | Computation of feedback value | Normalized difference between average activity in the SAN minus the average DMN activity |
| | Target ROI | 4 SAN ROIs and 4 DMN ROIs |
| | Crossover design? | No |
| | Up/downregulation | Upregulation only (SAN > DMN) |
| | Stimulus during regulation block | Black up-arrow |
| | Stimulus during baseline block | Black square |

| | | |
|---|---|---|
| **Amano et al., 2016** | General instruction for regulation blocks | "Somehow regulate their brain activity during the presentation of the achromatic grating to make the size of the solid gray disk presented during the subsequent feedback period as large as possible" |
| | General instruction for baseline | "Maintain fixation on the fixation point for the whole experiment period" |
| | Number of training days | 3 |
| | Number of training runs | Maximum of 12 runs |
| | Duration of the training run | 6min30s |
| | Number of baseline blocks | 15 + 1 in the beginning |
| | Duration of baseline block | initial baseline 30 s, fixation 7 s, ISI 6 s |
| | Number of regulation blocks | 15 |
| | Duration of regulation block | 6 s |
| | Another condition during training run | No |
| | Feedback representation | Disk |
| | Duration of feedback block | 1 s |
| | Computation of feedback value | "Multiplying the BOLD-signal multi-voxel patterns with the weights determined in the color classifier construction stage, and by passing this weighted sum through a logistic function (likelihood estimates could range from 0 to 100%)" |
| | Target ROI | Multivoxel patterns in V1/V2 |
| | Crossover design? | No |
| | Up/downregulation | Upregulation |
| | Stimulus during regulation block | "An achromatic (gray black) vertical grating, identical to the one used in the color classifier construction stage presented with a fixation point at the center of the screen" |
| | Stimulus during baseline block | Fixation point on a black background |
| **Scheinost et al., 2020** | General instruction for regulation blocks | "Keep the gauge as close to full as possible", subjects were informed that it was related to attention |
| | General instruction for baseline | No |
| | Number of training days | 1 |
| | Number of training runs | 3 |
| | Duration of the training run | 14 min |
| | Number of baseline blocks | 0 |
| | Duration of baseline block | 0 |
| | Number of regulation blocks | 4 |
| | Duration of regulation block | 3 min |
| | Another condition during training run | No |
| | Feedback representation | Gas gauge |
| | Duration of feedback block | 30 s |
| | Computation of feedback value | Normalized difference between the mean strength of functional connections in the high minus low attention networks |
| | Target ROI | High and low attention networks |
| | Crossover design? | No |
| | Up/downregulation | Gas gauge to the right |
| | Stimulus during regulation block | gradual-onset continuous performance task |
| | Stimulus during baseline block | No |
| **Krause et al., 2021** | General instruction for regulation blocks | "Either increase or decrease the size of the disc on the screen with their brain ... They were told that they could achieve this by thinking of something specific, performing some mental task internally, or getting into a certain mood, emotion, feeling, or state of mind, and that they had to explore different mental strategies to find one that works for them" |
| | General instruction for baseline | Not informed |
| | Number of training days | 3 |
| | Number of training runs | 7 to 8 |
| | Duration of the training run | 10 min |
| | Number of baseline blocks | 1 |
| | Duration of baseline block | 34 s in the beginning, 10 s in the subsequent blocks |
| | Number of regulation blocks | 16 (8 upregulation and 8 downregulation) |
| | Duration of regulation block | 16 s |
| | Another condition during training run | Yes, delay for 6 s |
| | Feedback representation | Gray disc superimposed with a black circle with radius varying depending on feedback. Orange dot in the center |
| | Duration of feedback block | 4 s |
| | Computation of feedback value | Feedback was based on the difference signal between the averages (mean) of all voxels in the SN and ECN ROIs, with the direction being alternated between participants (participants 1, 3, 5, 7, 9, 11: SN - ECN; participants 2, 4, 6, 8, 10: ECN - SN). |
| | Target ROI | SN and ECN ROIs |
| | Crossover design? | No |
| | Up/downregulation | Yes. In the first and second run of the first Training session, all regulation blocks were of the condition “larger” and “smaller”, respectively |
| | Stimulus during regulation block | Disc of radius half of the maximum size with outward or inward arrows to represent up- and down-regulation, respectively. Green dot in the center |
| | Stimulus during baseline block | Disc of radius half of the maximum size. Black dot in the center |

#### 4.3. Table S3

*Table S3. Significant clusters in whole-brain analyses for Model 1 (Fig. 1A), using a threshold set at a voxel-wise uncorrected p < 0.001 and cluster-wise FDR-corrected p < 0.05. Regions were automatically labeled using the AnatomyToolbox atlas. Local maxima are reported when separated by more than 10 mm and a maximum of ten peaks per cluster was set.*

| Region Label | Extent (voxels) | t-value | MNI Coordinates | | |
|---|---|---|---|---|---|
| | | | x | y | z |
| L Olfactory cortex | 30105 | 7.015 | -12 | 2 | -8 |
| R Caudate Nucleus | | 6.779 | 12 | 6 | -4 |
| L Mid Orbital Gyrus | | 6.386 | -4 | 44 | -6 |
| R Cerebellum (Crus 1) | | 6.042 | 36 | -72 | -30 |
| R Mid Orbital Gyrus | | 5.932 | 8 | 40 | -6 |
| L Mid Orbital Gyrus | | 5.664 | -6 | 34 | -8 |
| R Rectal Gyrus | | 5.591 | 6 | 48 | -12 |
| R Cerebellum (Crus 2) | | 5.379 | 12 | -80 | -30 |
| L IFG (p. Orbitalis) | | 5.366 | -24 | 20 | -20 |
| R IFG (p. Orbitalis) | | 5.248 | 30 | 22 | -4 |
| L Angular Gyrus | 2662 | 5.168 | -40 | -72 | 42 |
| L Angular Gyrus | | 5.059 | -46 | -64 | 34 |
| Location not in atlas | | 4.686 | -36 | -60 | 20 |
| L Inferior Parietal Lobule | | 4.503 | -50 | -58 | 52 |
| L Inferior Parietal Lobule | | 4.500 | -34 | -74 | 52 |
| L Inferior Parietal Lobule | | 3.983 | -52 | -38 | 42 |
| L Inferior Parietal Lobule | | 3.923 | -34 | -58 | 50 |
| L Inferior Parietal Lobule | | 3.907 | -56 | -44 | 54 |
| L Superior Parietal Lobule | | 3.714 | -20 | -80 | 54 |
| L Superior Parietal Lobule | | 3.353 | -18 | -74 | 62 |

#### 4.4. Table S4

*Table S4. Significant clusters in whole-brain analyses for Model 2 (Fig. S1, association between connectivity with the nucleus accumbens and feedback scores). Regions were labeled based on the meta-analytic associations of Neurosynth.*

| Region Label | Extent (voxels) | t-value | MNI Coordinates | | |
|---|---|---|---|---|---|
| | | | x | y | z |
| R superior parietal lobule | 43 | 4.032 | 50 | -42 | 62 |
| Substantia nigra | 107 | 3.948 | 12 | -18 | -14 |
| R anterior insula | 12 | 3.326 | 40 | 18 | -10 |

#### 4.5. Table S5

*Table S5. Clusters of a thresholded effect-size map in whole-brain analyses for Model 3 (Fig. S2) (|t| > 2.46). Regions were automatically labeled using the AnatomyToolbox atlas. Local maxima are reported when separated by more than 10 mm and a maximum of ten peaks per cluster was set.*

| Region Label | Extent (voxels) | t-value | MNI Coordinates | | |
|---|---|---|---|---|---|
| | | | x | y | z |
| R Superior Temporal Gyrus | 203 | 3.185 | 62 | -34 | 8 |
| R Posterior-Medial Frontal | 182 | 3.945 | 6 | 6 | 80 |
| Location not in atlas | 108 | 3.422 | 16 | 18 | 28 |
| L Paracentral Lobule | 52 | 3.272 | -14 | -26 | 82 |
| Location not in atlas | 51 | 3.180 | 34 | 24 | 18 |
| Location not in atlas | 49 | 3.716 | -68 | -40 | 18 |
| Location not in atlas | 48 | 3.278 | -18 | 22 | -6 |
| R Precentral Gyrus | 47 | 3.693 | 14 | -32 | 84 |
| Location not in atlas | 23 | 3.395 | 30 | 54 | -18 |
| Location not in atlas | 23 | 3.164 | -12 | -40 | 20 |
| R Superior Temporal Gyrus | 15 | 2.767 | 66 | -36 | 28 |
| R SupraMarginal Gyrus | 12 | 2.935 | 68 | -24 | 34 |
| Location not in atlas | 11 | 2.880 | 28 | -26 | 26 |
| Location not in atlas | 10 | 2.724 | -12 | -18 | -36 |
| L Precentral Gyrus | 7 | 2.711 | -48 | 2 | 52 |
| Location not in atlas | 6 | 2.740 | 28 | -34 | 16 |
| Location not in atlas | 256 | -4.555 | -18 | 18 | -26 |
| L Inferior Parietal Lobule | 81 | -3.363 | -34 | -72 | 54 |
| R Superior Occipital Gyrus | 47 | -2.783 | 24 | -88 | 34 |
| L Superior Frontal Gyrus | 29 | -2.983 | -20 | 52 | 42 |
| Location not in atlas | 28 | -3.275 | 26 | 32 | -24 |
| Location not in atlas | 26 | -3.391 | -6 | 36 | 64 |
| L Middle Frontal Gyrus | 26 | -3.126 | -42 | 26 | 50 |
| R Superior Frontal Gyrus | 24 | -3.024 | 20 | 30 | 64 |
| L Inferior Parietal Lobule | 24 | -2.900 | -44 | -56 | 60 |
| Location not in atlas | 20 | -3.074 | -36 | 16 | -38 |
| L Calcarine Gyrus | 17 | -2.752 | -4 | -100 | 12 |
| L Superior Medial Gyrus | 17 | -2.586 | -8 | 64 | 12 |
| Location not in atlas | 16 | -2.941 | -36 | 60 | 16 |
| Location not in atlas | 15 | -3.001 | 24 | -10 | -34 |
| L Cuneus | 15 | -2.825 | -6 | -92 | 38 |
| L Superior Medial Gyrus | 9 | -2.675 | -6 | 48 | 54 |
| Location not in atlas | 8 | -2.975 | -6 | 56 | 48 |
| L Fusiform Gyrus | 6 | -2.683 | -38 | -36 | -16 |
| Location not in atlas | 6 | -2.642 | 54 | -42 | -24 |
| Location not in atlas | 5 | -2.774 | 2 | 44 | -30 |

| | | | | | |
|---|---|---|---|---|---|
| R Superior Occipital Gyrus | 5 | -2.553 | 16 | -96 | 24 |
| Location not in atlas | 5 | -2.532 | -48 | 36 | -18 |

#### 4.6. Table S6

*Table S6. Clusters of a thresholded effect-size map in whole-brain analyses for Model 4 (Fig. S3) (|t| > 2.38). Regions were automatically labeled using the AnatomyToolbox atlas. Local maxima are reported when separated by more than 10 mm and a maximum of ten peaks per cluster was set.*

| Region Label | Extent (voxels) | t-value | MNI Coordinates | | |
|---|---|---|---|---|---|
| | | | x | y | z |
| L Cerebelum (Crus 2) | 196 | 3.860 | -34 | -84 | -34 |
| L Cerebelum (Crus 1) | 196 | 3.794 | -48 | -80 | -20 |
| L Orbitofrontal cortex | 131 | 3.302 | -40 | 46 | -12 |
| L Cerebelum (Crus 1) | 20 | 2.561 | -22 | -90 | -20 |
| Location not in atlas | 16 | 2.712 | 38 | 62 | 12 |
| Location not in atlas | 8 | 2.847 | -20 | 22 | -26 |
| Location not in atlas | 7 | 2.651 | -30 | 40 | -18 |
| L Rectal Gyrus | 6 | 2.664 | -4 | 8 | -16 |
| L Cerebelum (VI) | 61 | -3.180 | -40 | -40 | -28 |
| Location not in atlas | 51 | -2.973 | 4 | -40 | -20 |
| L Middle Frontal Gyrus | 26 | -2.868 | -22 | 22 | 64 |
| R Postcentral Gyrus | 23 | -2.786 | 24 | -38 | 78 |
| R Precentral Gyrus | 22 | -2.774 | 50 | 4 | 52 |
| R Superior Frontal Gyrus | 12 | -2.613 | 20 | -16 | 80 |
| L Precentral Gyrus | 8 | -2.670 | -18 | -22 | 82 |
| L Superior Temporal Gyrus | 7 | -2.985 | -68 | -16 | 8 |
| R ParaHippocampal Gyrus | 6 | -2.456 | 30 | -30 | -20 |
| Location not in atlas | 5 | -2.645 | -10 | -14 | -22 |
